# Monosynaptic connections link functionally similar regions in human cortex

**DOI:** 10.64898/2026.05.19.726279

**Authors:** Rui Xu, Alan Bush, Atsushi Takahashi, Nathaniel Sisterson, Ashley Walton, Samuel Hutchinson, Alexandra Kammen, Clemens Neudorfer, Vasileios Kokkinos, Pranav Nanda, Niharika Jhingan, Ammar Marvi, Alexander Fung, Suseendrakumar Duraivel, Eghbal Hosseini, Colton Casto, Claudia Valenzuela, Seonghwan Yee, John Kirsch, Evelina Fedorenko, Nancy G. Kanwisher, Robert Desimone, R. Mark Richardson

## Abstract

The functional organization of human cortex has been mapped in considerable detail^1–5^, yet the axonal connections linking these areas remain largely unknown. This gap precludes inferring either the computational pathways through cortical networks or the principles governing why each area connects to its particular target regions. Here, in patients undergoing intracranial monitoring for epilepsy, we used concurrent electrical stimulation and functional magnetic resonance imaging (es-fMRI), validated previously against tracer studies in nonhuman primates^6^, to map anatomical connectivity of cortical sites across the whole brain, and combined these maps with task and resting-state fMRI in the same individuals. Es-fMRI revealed four main findings. First, connectivity followed an asymmetric functional-similarity principle: connected sites tended to share similar functional profiles, but, because connectivity is sparse, most pairs of functionally similar sites were not connected. Second, although connectivity also declines with distance^7^, the functional profile explained three times as much variance in connectivity as distance did, and long-range connections were the most functionally specific. Third, es-fMRI revealed direct monosynaptic links between established functional regions, including between the fusiform face area (FFA)^2^ and the temporoparietal junction (TPJ)^3^ for social cognition. Fourth, resting-state functional connectivity (rsFC) of a stimulation site was only weakly correlated with that site’s es-fMRI connectivity. Together, these results provide a first principled link between anatomical connectivity and functional organization in the human cortex, and establish es-fMRI as a scalable approach to building a tracer-grade connectome of the human brain.

## INTRODUCTION

Decades of neuroimaging research have mapped the functional organization of the human cerebral cortex in considerable detail. Functionally specialized regions for face and object recognition, language, theory of mind, executive control, and other cognitive functions have been identified at consistent cortical locations across virtually all neurotypical adults^1–5^, and their causal importance has been established by selective deficits in patients with focal lesions and by behavioral and perceptual changes induced by direct cortical stimulation^8–11^. The regions supporting higher-level cognitive functions, such as language or executive control, are often described as forming large-scale brain ‘networks’, whose constituent regions share response profiles and exhibit synchronized spontaneous activity^12–16^. Yet, existing methods in humans reveal little about how information flows between regions, which requires knowledge of *anatomical connectivity,* the axonal connections between regions. Without such knowledge, we cannot characterize the computations within a region, the interactions between regions, the larger network each region inhabits, or the general principles that explain why some regions connect to particular others.

Tracer studies in nonhuman primates (NHPs), now spanning decades and hundreds of injections, have revealed that cortical connectivity is extraordinarily complex yet governed by several organizing principles, including hierarchical organization^17,18^, topographic mappings^6,19,20^, distance dependence^7,21^, and laminar pattern dependence^22^. NHPs, however, cannot produce maps of certain higher cognitive functions, such as theory of mind and language, that define much of human cortical organization. As a result, in the absence of methods for unambiguously establishing anatomical connections in humans, the relationship between anatomical connectivity and cortical function in the human brain remains largely unknown.

Every existing method for probing human cortical connectivity has well-documented limitations. Diffusion MRI-based tractography (dMRI-tractography)^23–27^ is the most widely used, yet it suffers from inherent limitations: superficial white matter fiber systems impede detection of long-range cortical connections^28^, and the tractography outcome correlates poorly with gold-standard tracer results^6,29^. A recent approach^30,31^ constrains dMRI-tractography with NHP tract tracing, but it relies on NHP-human homologies^32,33^ and is restricted to short-range connections. Resting-state functional connectivity (rsFC)^34^ is another common surrogate, but it reflects the co-fluctuation of spontaneous neural activity rather than direct anatomical connections^35^. Cortico-cortical evoked potentials (CCEPs) recorded during direct cortical stimulation in epilepsy patients have provided valuable insights into signal propagation across human brain networks^36–41^, but electrode coverage is sparse, and the technique has not been systematically validated against anatomical tracers.

Here, we used concurrent electrical stimulation and whole-brain fMRI (es-fMRI) in epilepsy patients to map anatomical connectivity of individual cortical sites across the entire brain, and combined it with task-based fMRI and resting-state fMRI in the same individuals. Es-fMRI has been validated in NHPs as a close surrogate for monosynaptic tracer-based connectivity^6,20,42–44^, and is, to our knowledge, the only method in humans that can resolve whole-brain connectivity at the level of monosynaptic projections. Our multi-modal human data reveal an asymmetric functional-similarity principle: a stimulation site’s connections preferentially target regions that share its functional profile, but, because connectivity is sparse, many pairs of functionally similar regions lack a direct connection. The same data also yield an initial sketch of the structural connections of established functional regions such as the fusiform face area, language regions, and the temporoparietal junction (TPJ) for social cognition, providing a foundation for building a tracer-grade map of the human cortex.

## RESULTS

### Conducting es-fMRI in humans

The primary safety risk in es-fMRI is tissue heating^45–47^. Through extensive phantom tests (see METHODS), we identified the distance between the extension cables and the transmit coil as a key, but previously underappreciated factor in tissue heating. With optimal cable positioning ensured by a custom-made track, we recorded insignificant^48^ heating in all tests using es-fMRI sequences (<0.5 °C; e.g., Extended Data Fig. 1b, T1w/EPI). Because high-SAR sequences combined with poor cable placement could lead to severe heating (>10 °C within 2 minutes), we strongly recommend safety testing under the specific conditions to be used before initiating es-fMRI.

We combined these cable positioning methods with state-of-the-art MRI coils and sequences in es-fMRI to optimize signal, along with stimulation parameters previously validated in NHP studies^6,42^ (see METHODS). To reduce MRI artifacts (Extended Data Fig. 1c) and further limit heating risk, we removed most stereoelectroencephalography (SEEG) electrodes at the bedside after epilepsy monitoring had concluded, retaining a few (typically 2-4) containing planned stimulation sites for es-fMRI, which were removed shortly afterward. Bedside removal of SEEG electrodes is an established clinical procedure, and our experimental considerations affected only the order of electrode removal.

The effective signal-to-noise ratio (SNR), inferred from t-statistics normalized to equal scan time (see METHODS), exceeded that of the only previously published human es-fMRI dataset^49^ by more than 4-fold and matched our NHP es-fMRI study that used contrast agents^6^ (Fig. 1a). The SNR of es-fMRI was also more than twice that of task fMRI acquired with similar scanning parameters (see METHODS), highlighting the efficacy of intracranial stimulation in driving fMRI signal.

**Figure 1.**
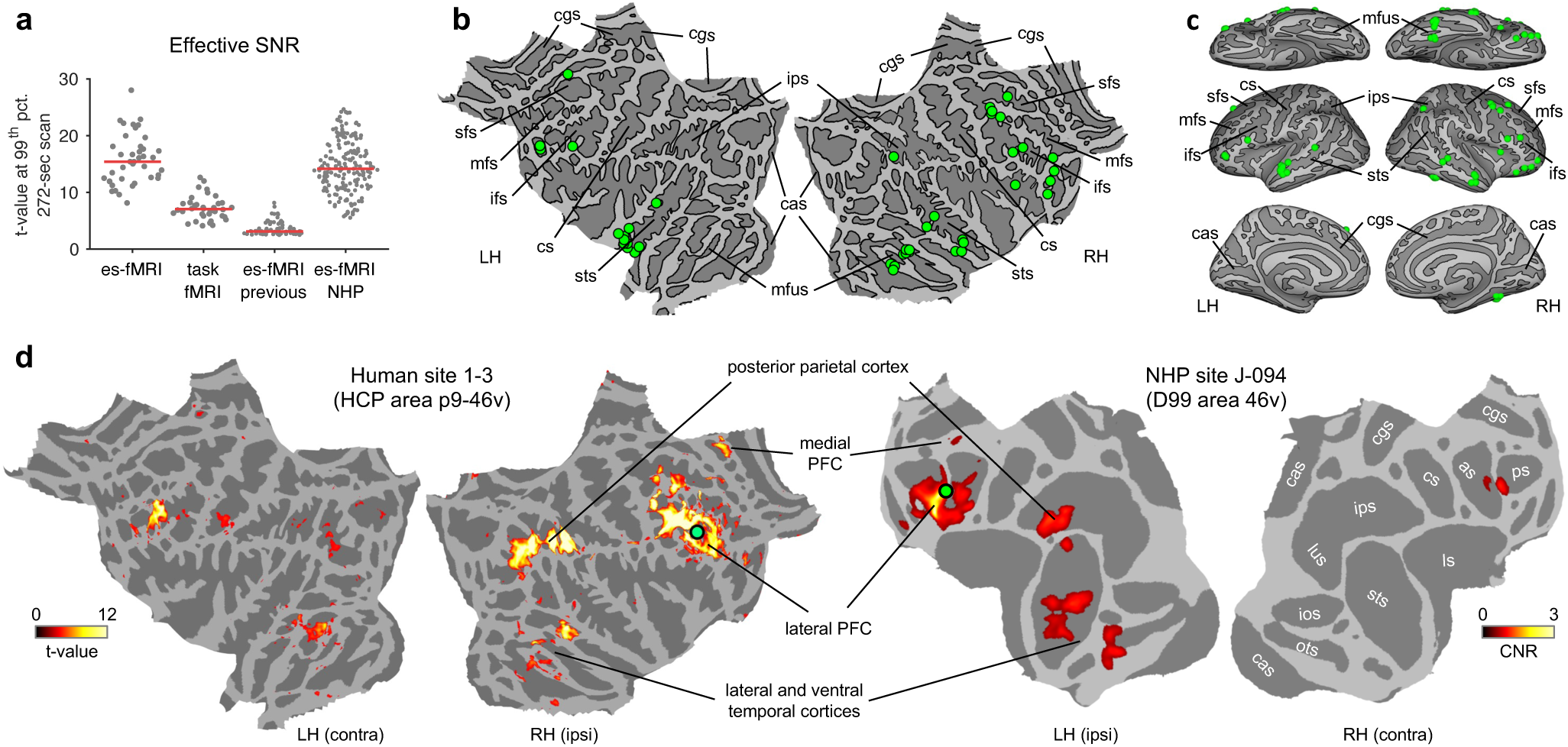
Conducting es-fMRI in humans. (a) Effective signal-to-noise ratio (SNR) comparison across studies, species, and fMRI modalities. Each dot represents a stimulation site or task contrast from one subject. Effective SNR is quantified as the t-statistic at the 99th percentile of significantly activated voxels, normalized to a 272-second scan. Datasets compared: our es-fMRI (human), our task fMRI (human), the one previous human es-fMRI study^49^, and our NHP es-fMRI study^6^. (b) Locations of 42 stimulation sites projected onto flattened cortical surfaces. LH/RH, left/right hemisphere; cas, calcarine s. (sulcus); cgs, cingulate s.; cs, central s.; ifs, inferior frontal sulcus; ips, intraparietal s.; mfs, middle frontal sulcus; mfus, mid-fusiform s.; sfs, superior frontal s.; sts, superior temporal s. (c) Stimulation site locations on inflated cortical surfaces. (d) Cross-species comparison of es-fMRI connectivity patterns at matching cortical locations. The left panel shows human PBRs on flattened surfaces (left hemisphere-contralateral, right hemisphere-ipsilateral), and the right panel shows NHP PBRs (left-ipsilateral, right-contralateral). Human and NHP sites were matched based on sulcal patterns and area labels. HCP, hman connectome project; as, arcuate s. (sulcus); ios, inferior occipital s.; ls, lateral s.; lus, lunate s.; ots, occipitotemporal s.; ps, principal s.

We stimulated 42 association-cortical sites in eight participants (three males, five females; age=39±11 yrs), predominantly in the prefrontal and temporal cortices (Fig. 1b,c). We measured positive BOLD responses (PBRs) to electrical stimulation because studies in NHPs have shown that PBRs from cortical stimulation are highly correlated with monosynaptic anterograde connections in the cortex, whereas negative BOLD responses (NBRs) reflect polysynaptic propagation^43^. Our previous results from an es-fMRI study in macaque lateral prefrontal cortex (LPFC) were consistent with this interpretation. Because anterograde and retrograde connections from tracer studies in animals are largely reciprocal, we interpret the PBRs as reflecting mainly bidirectional connectivity and, for simplicity, refer to them as “connections”.

Although our electrode coverage in LPFC was too sparse to perform a quantitative cross-species comparison with our NHP study, individual examples from LPFC showed remarkable cross-species similarity. Figures 1d and Extended Data Fig. 2 show significant stimulation-elicited PBRs of three pairs of human and NHP sites spaced dorsally-to-ventrally in the LPFC. At approximately matching locations, the spatial patterns of ipsilateral PBRs appeared highly similar between species. Extended Data Fig. 3a shows the full connectivity matrix between all 42 stimulation sites (see Extended Data Fig. 3b for the location of each) and ipsilateral HCP areas.

Across all human sites, we found far more extensive PBRs in the ipsilateral than in the contralateral cortex, consistent with our NHP es-fMRI data^6^ and tracer studies. The sparseness of ipsilateral connections was also similar across species, quantified by the percentage of surface area showing PBRs per site (humans: ipsi, 6.7±3.9%; contra, 1.0±1.6%; ipsi vs. contra, t_41_=11.4, p=3.0×10^−14^. NHPs: ipsi, 8.0±5.5%; contra, 2.1±2.0%; t_174_=18.9, p=5.2×10^−44^). The strong ipsilateral preference is consistent with the interpretation that cortical PBRs of es-fMRI correspond to monosynaptic connections^42,43^. As in NHPs, PBRs in humans were typically confined to a few “clusters” rather than scattered throughout the cortex.

### Connectivity is functionally specific

To relate connectivity and function, we recruited the same participants for noninvasive MRI sessions when they were not implanted with SEEG electrodes. We conducted well-established task-based fMRI localizers (Extended Data Table 1) that produce replicable activations at the individual level in healthy subjects. Together, the localizers comprised 32 contrasts spanning 13 functions across high-level visual processing (face, body, scene, and object perception)^1,2,50,51,^ high-level auditory processing (speech perception)^52^, and several aspects of cognition (executive control supported by the multiple demand network or MD, intuitive physical reasoning, language, theory of mind or TOM, episodic memory and prospection or episodic projection, and functions supported by the default mode network or DMN)^3–5,53–56^. The fMRI activation patterns in our participants (Extended Data Fig. 4) were broadly consistent with prior results in neurotypical adults^57,58^. Multiple contrasts for the same function were included to improve functional mapping reliability, yielding activation patterns that were highly similar, as expected (Extended Data Fig. 5). We tailored the selection of localizers to participants’ electrode coverage and available time (Extended Data Table 2), resulting in 22.1±6.0 contrasts and 11.1±0.8 functions per participant.

Our data thus enable us to ask, for any given cortical site, how its anatomical connectivity (revealed by es-fMRI) relates to functional response profiles across the brain (revealed by multiple task fMRI activations in the same individual). A first look at this relationship suggests a principle we will call asymmetric functional similarity: the connections of a stimulation site are most common to areas that share a similar functional profile, but many areas with a similar functional profile do not have a connection with the stimulation site, reflecting sparse connectivity. For example, Figure 1d shows that stimulation site 1-3 in the right LPFC is anatomically connected to regions in ipsilateral prefrontal, parietal, and temporal cortices. This site was most strongly activated in an MD contrast (MD.Math), and most of its connected regions, within and outside LPFC, also showed activations in the same contrast, while largely avoiding activations in the language, TOM, DMN, and object perception contrasts (Extended Data Fig. 6a). Yet many other MD-responsive regions in the same individual showed no connection to site 1-3.

Another stimulation site (site 1-1; Extended Data Fig. 6b) in the left anterior superior temporal sulcus (STS) showed anatomical connections to many temporal regions at similar dorsal-ventral levels, as well as scattered prefrontal and posteromedial regions. This site was most strongly activated in a language contrast (Lang.NWAud), and most of its connected regions also showed activations in the same contrast, although some language-responsive regions remained unconnected.

To test this principle across the population of stimulation sites, we introduced a metric of functional similarity (see METHODS), defined as the cosine similarity between the functional profile (a vector of task fMRI activations across all functions) of a stimulation site (averaged across nearby voxels, i.e., ≤4 mm) and that of each non-adjacent voxel. The metric’s value ranges from 0 (indicating no overlap in the task activations between the stimulation site and a non-adjacent voxel) to 1 (indicating identical functional profiles). We marked a voxel as unknown if it showed no task activation (and thus no overlap with the functional profile of the stimulation sites). Figures 2a and 2c show the functional similarity of all connected voxels for sites 1-3 and 1-1, respectively. Consistent with Extended Data Fig. 5, the majority of the connected voxels had functional profiles very similar to those of the respective sites, with a small portion showing low or unknown similarity. Conversely, the majority of the unconnected voxels exhibited low or unknown similarity (Fig. 2b,d).

**Figure 2.**
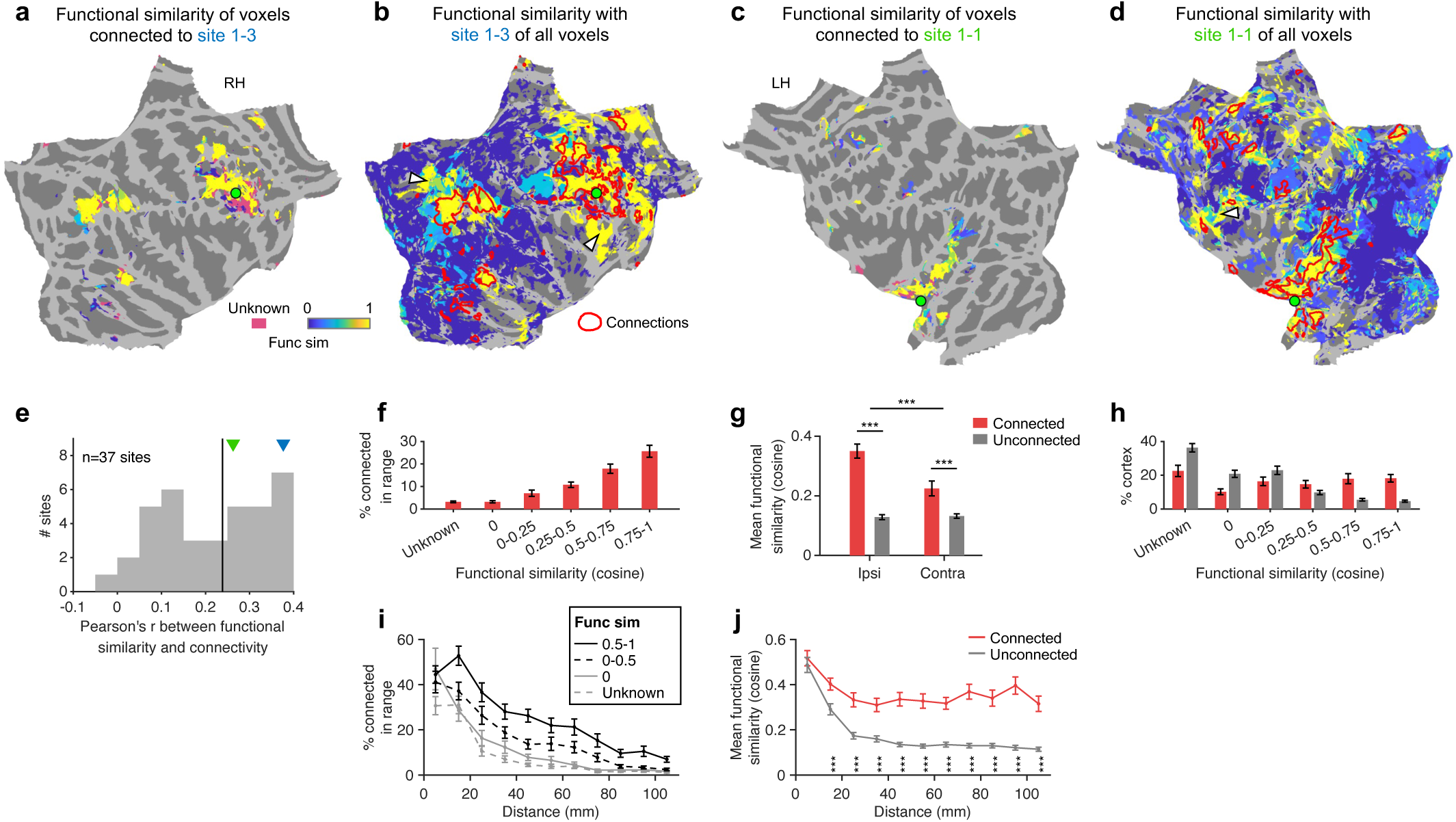
Connectivity is functionally specific. (a) Functional similarity map for ipsilaterally connected voxels (shown in Fig. 1d) of example stimulation site 1-3 (a), displayed on the flattened cortical surface. Color indicates cosine similarity between the functional profile of the stimulation site (averaged across nearby voxels) and each connected voxel, ranging from 0 (no shared functions) to 1 (identical profiles). Pink indicates unknown similarity. (b) Functional similarity maps for all voxels of site 1-3. Red contours encircle connected voxels. White arrows indicate unconnected voxels with high functional similarity. Voxels with unknown functional similarity were not colored. (c,d) Same as (a,b) but for stimulation site 1-1. (e) Histogram across 37 functionally defined stimulation sites of the Pearson correlation between functional similarity and connectivity of a given stimulation site to all ipsilateral voxels that are not near the site. Vertical line indicates the median. Colored triangles mark the example sites. (f) Percentage of connected voxels across functional similarity ranges. (g) Mean functional similarity of connected and unconnected voxels, separately for the ipsilateral and contralateral hemispheres. Unknown similarities were set to zero. All other panels show ipsilateral results only. (h) Distribution of functional similarity for connected (red) and unconnected (gray) voxels. (i) Percentage of connected voxels as a function of distance from stimulation sites, plotted separately for different functional similarity ranges. (j) Mean functional similarity of connected and unconnected voxels as a function of distance. Unknown similarities are set to zero. All error bars denote SEM across stimulation sites. ***: p<0.001.

Despite the strong link between connectivity (thresholded and truncated t-value; see METHODS) and functional similarity (cosine) in the example sites, the correlation between them was only moderate (Pearson’s r: site 1-3, 0.38; site 1-1, 0.26). Site 1-1 had a correlation that was about the same as the median across 37 functionally defined sites (Fig. 2e; median/mean/std=0.24/0.22/0.12), and site 1-3 was at the higher end of the distribution. Stimulation sites with too few nearby voxels activated in any contrast were considered functionally undefined (n=5; see METHODS). The correlation values were affected by the sparse connectivity: only a small portion of voxels were connected even when their functional similarity was at its highest level (Fig. 2f; 25.6%, 0.75<similarity≤1). White arrows in figures 2b and 2d highlight some unconnected voxels with high similarity for the example sites (see Extended Data Fig. 7a,c for sparseness across similarity levels for these sites).

We next examined these relationships across all 37 functionally defined stimulation sites (Fig. 2f-j). The functional specificity of the observed connections is evident in the increasing sparsity at lower levels of functional similarity (Fig. 2f; e.g., 3.20/3.22% at unknown/zero similarity). As an alternative measure, we compared the mean functional similarity between connected and unconnected voxels (Fig. 2g), with unknown similarities set to zero. Connected voxels were significantly more functionally similar than unconnected ones in both hemispheres (ipsi: t_36_=11.1, p=4.0×10^−13^; contra: t_36_=4.0, p=2.8×10^−4^), and the difference was significantly greater in the ipsilateral cortex (t_36_=4.8, p=3.2×10^−5^), suggesting that contralateral connections are less functionally specific. Figure 2h shows the distribution of functional similarity (not just the mean) for ipsilaterally connected and unconnected voxels (see Extended Data Fig. 7b,d for distribution at the example sites). A larger portion of connected voxels fell within higher similarity ranges (>0.25), while fewer fell in the lower ranges, compared to unconnected voxels. However, it is worth noting that a non-trivial percentage of connected voxels showed no functional overlap with stimulation sites (22.6/10.2% at unknown/zero similarity). These could represent cross-network connections, which are considered further below.

Within each level of functional similarity, connectivity became increasingly sparse at longer distances (Fig. 2i), consistent with the well-known connectivity-distance relationship in NHPs^7^. Functional specificity of connections was preserved across distances and became more pronounced at greater distances: the difference in mean functional similarity between connected and unconnected voxels was smaller at close distances and increased at longer distances (Fig. 2j). Thus, long-range connections are sparse but particularly functionally specific.

### Functional profile predicts connectivity

The functional similarity analysis above used a single composite metric with equal weights across functions. A natural next question is whether task fMRI activation patterns for certain functions are more strongly associated with connectivity than others. To address this, we first examined activations across individual contrasts, separately, to assess how each was correlated with connectivity. Figure 3a shows the percentage of voxels near each of the 37 functionally defined stimulation sites (rows) activated in each of the 32 task fMRI contrasts (columns, with labels at the top and bottom of the matrix). We ordered the columns and rows to make the quasi-diagonal pattern apparent, highlighting that each site was typically activated in a subset of task-derived functions/contrasts. Figure 3b shows the corresponding matrix with the correlation (Pearson’s r) between the cortex-wide connectivity pattern of each stimulation site, and the activation pattern of each contrast, with the same ordering as Fig. 3a. Thus, for example, the first row of Fig. 3a shows that stimulation site 6-2 has high activations in the object perception contrasts, and the first row of Fig. 3b shows that its connectivity pattern also had a high correlation with the activation patterns of the object perception contrasts. The correlation matrix (Fig. 3b) exhibited a diagonal pattern strikingly similar to that in Fig. 3a, indicating that connectivity-activation correlations tended to be positive when a stimulation site (row) was activated in a contrast (column), and low or even negative when it was not.

**Figure 3.**
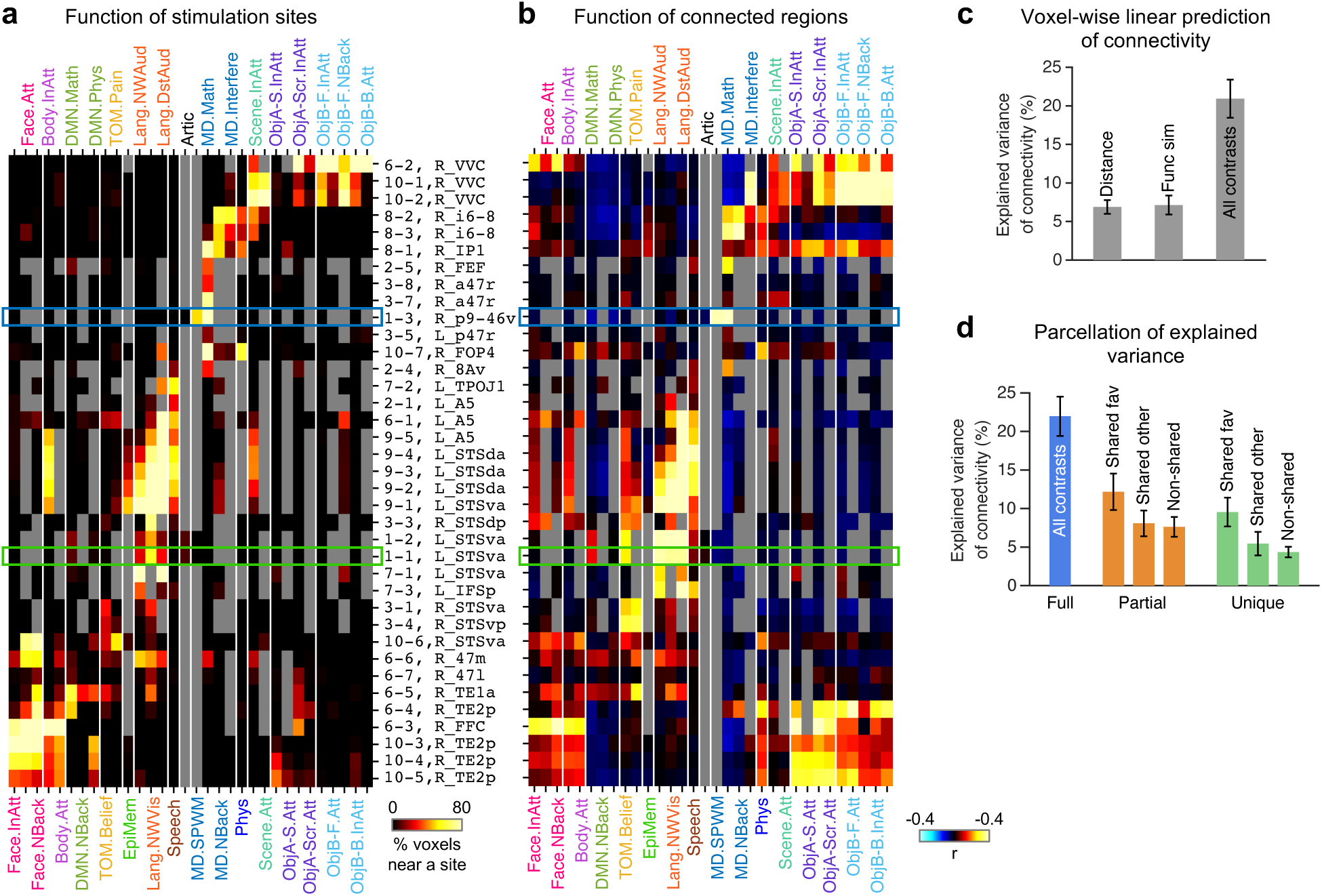
Functional profile predicts connectivity. (a) Contrast×site matrix showing the percentage of voxels near each of 37 functionally defined stimulation sites (rows) activated in each of 32 task fMRI contrasts (columns). Rows and columns are ordered to highlight the quasi-diagonal pattern. Gray entries indicate missing data (contrasts unavailable for a participant). Colored rectangles mark the example sites in Fig. 2. (b) Contrast×site matrix showing the Pearson correlation between the cortex-wide connectivity pattern (excluding voxels near the site) of each site and the activation pattern of each contrast, with the same ordering as (a). (c) Cross-validated variance explained (R^2^) by linear regression models predicting voxel-wise connectivity from different predictors. (d) Variance decomposition of three groups of contrasts defined based on each site’s functional profile. Each partial model used a different subset of contrasts. Fav: favorite. All error bars denote SEM across sites.

This similarity could be trivial if it merely reflected the general tendency for nearby brain regions to be both connected and functionally similar^12,59^. To rule out this possibility, we used SPIN permutation^60^ to jointly rotate and displace all stimulation sites and their connections on the cortical surface while keeping task activations fixed. This operation disrupted the correspondence between connectivity and function but preserved their shared dependence on distance and their within-modality covariance. The observed similarity between matrices (Spearman’s ρ) was higher than all permuted values (Extended Data Fig. 8, top; actual=0.579, permuted median/max=0.198/0.575). We further tested the influence of shared distance dependence by repeating the analysis while excluding voxels <40 mm from the stimulation sites. The actual similarity was largely unchanged, whereas the permuted similarity approached zero (Extended Data Fig. 8, bottom; actual=0.532, permuted median/max=0.026/0.471).

We next examined all contrasts jointly by predicting voxel-wise connectivity from the contrasts with linear regressors, using leave-one-HCP-area-out cross-validation (see METHODS). Activations across all contrasts explained 20.9% of the variance in connectivity (Fig. 3c, All contrasts), roughly threefold the variance explained by distance (Fig. 3c, Distance, 6.9%). The cosine functional similarity metric (7.1%) performed moderately, likely because it assigns equal weights to all contrasts and underestimates how well task fMRI responses predict connectivity.

Using the regression analysis, we compared the importance of three groups of contrasts (Fig. 3d): those belonging to the function that activated the largest number of voxels near a site (Shared favorite), other contrasts that activated the site (Shared other), and contrasts that did not activate the site (Non-shared). Unique variance for each group was computed as the difference between the full model R² and the R² of a model excluding that group (see METHODS). Despite comprising the fewest contrasts as predictors, the Shared favorite group explained the most (9.5% unique variance; 2.4±1.0 contrasts), followed by Shared other (5.4%, 4.9±2.9 contrasts) and Non-shared (4.4%, 16.2±4.7 contrasts). On a per-contrast basis, each Shared favorite contrast contributed approximately 4.0% of unique variance, compared to 1.1% per Shared other and 0.27% per Non-shared contrast. Thus, although all contrasts contributed to connectivity prediction, those shared with, and especially favored by, stimulation sites played a disproportionately greater role, reinforcing what we learned from examining contrasts individually (Fig. 3a,b). In other words, connections between two cortical locations are mainly determined by activations in functions that they share, rather than by non-shared activations.

### Connectivity across known fROIs

The analyses so far have been agnostic about the specific cortical locations of functional activations. Numerous studies, however, have localized various functions to particular cortical locations identifiable in virtually all neurotypical subjects. Such functionally defined regions (typically referred to as functional regions of interest, or fROIs) include many well-characterized areas, such as the fusiform face area (FFA) in the ventral temporal cortex^2^. The anatomical connections of these fROIs to other regions are highly informative for understanding how each fits into larger brain networks supporting complex functions, and our data include es-fMRI results for many of these fROIs.

To gain an understanding of the connections of specific fROIs, we defined each fROI following standard practice^58^ as the intersection of a previously established population-based mask for that fROI, and each individual’s task-based activations for the corresponding function (see METHODS). As an example, Figure 4 shows two stimulation sites (green crosses) in the FFA. Sites 10-3 (Fig. 4a-c) and 6-3 (Fig. 4d-f) from two participants were both located in the right fusiform gyrus at nearly identical cortical locations. The functional profiles for the two stimulation sites are shown in Fig. 4b and 4e. Site 10-3 showed a more typical face-selective activation profile (Fig. 4b and Extended Data Fig. 11b), whereas site 6-3 showed additional response to bodies (Fig. 4e and Extended Data Fig. 11e), typical for the fusiform body area (FBA)^61^. Both sites were connected to many face-responsive regions throughout the ventral temporal, lateral temporal, and lateral prefrontal cortices (Extended Data Fig. 9a,b), which fall along the “diagonal” connections in Fig. 3b. Thus, the FFA tends to have connections to other face-responsive areas, as has been shown in primates^62,63^.

**Figure 4.**
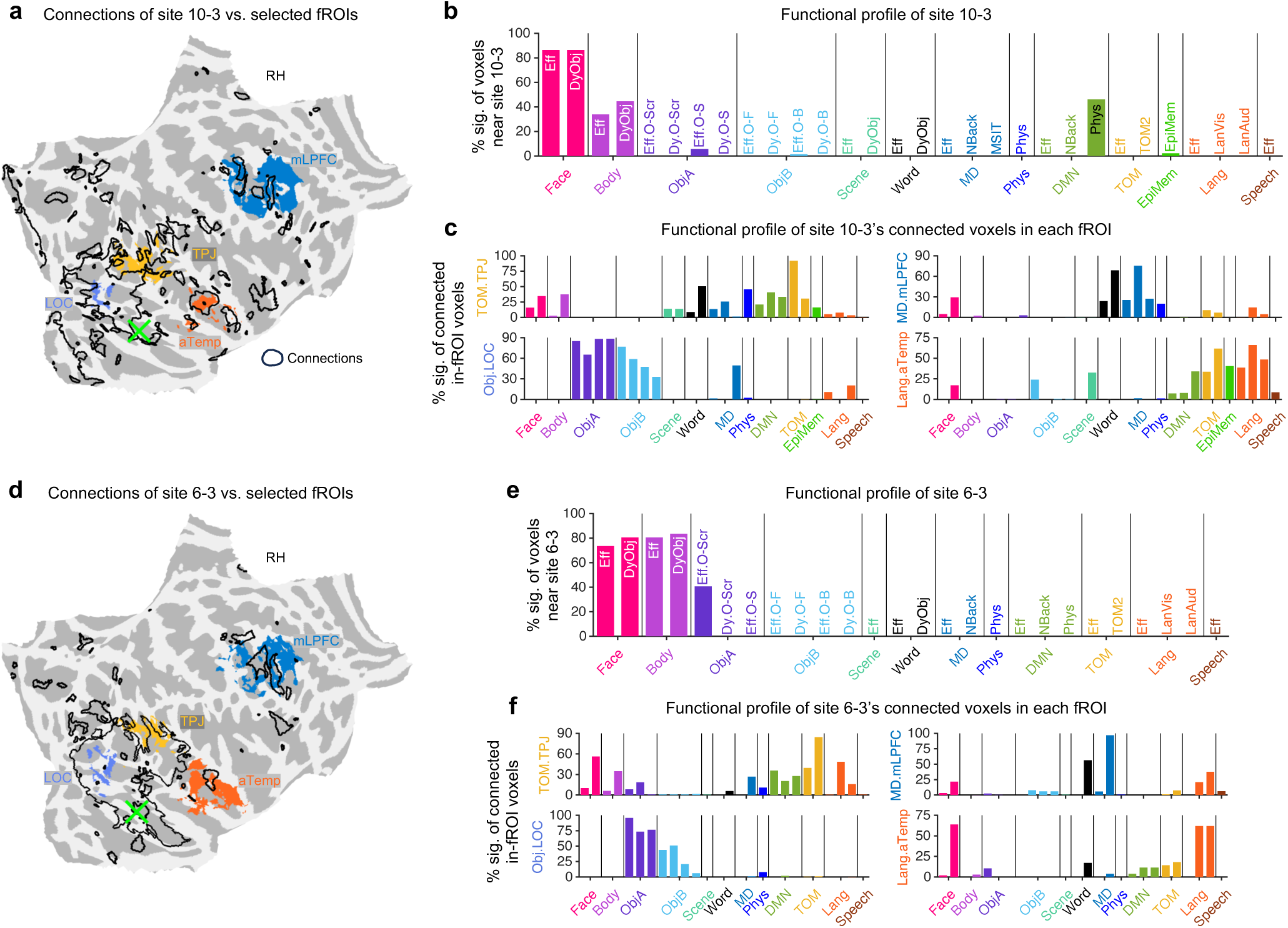
Connectivity of two FFA stimulation sites. (a-c) Results of stimulation site 10-3 (subject S10). (a) Site location (green X) on the right fusiform gyrus shown on the flattened cortical surface with fROIs as colored overlays and connections as contours. Colors of overlay match fROI names in (c). (b) Functional profile of site 10-3: percentage of voxels near the site activated in each contrast. Contrasts are sorted by function. Contrast labels indicate localizer abbreviation; ObjA and ObjB contrast labels additionally indicate control condition (e.g., Dy.O-Scr indicates objects > scrambled objects/words contrast of DyObj localizer). (c) Functional profile of connected voxels within each of four fROIs: percentage of connected voxels activated per contrast. Contrasts are in the same order as (b) (d-f) Same as (a-c) for site 6-3 (subject S6).

The FFA also showed “off-diagonal” connections to other areas with somewhat different functional profiles, which can give some insight into how the FFA interacts with other functional networks. For example, as shown in Figures 4a and 4d, both FFA sites were connected to some voxels within fROIs supporting object recognition (Obj.LOC), theory of mind (TOM.TPJ), executive control (MD.mLPFC), and language (Lang.aTemp). Figures S11A and S11D show only the connected voxels within fROIs. Task activations within these connected regions were not confined to a single function. We examined the functional profiles of the connected voxels within each fROI, shown as the percentage of voxels activated in each task-derived function (Fig. 4c,f). All such voxels in the TPJ showed activation in at least one TOM contrast by design, since the fROI was defined by TOM response. More informative were the additional responses observed there, e.g., those to faces, suggesting that these connections were driven, in part, by shared face responses, consistent with the finding that shared function predicts connectivity (Fig. 3). Face responses were also observed in the mLPFC MD fROI and in the aTemp language fROI, both of which had connections with the FFA. Extended Data Fig. 11 shows beta weights across conditions for stimulation sites and connected fROIs, broadly consistent with the percentage-based functional profiles in Fig. 4. Both FFA sites were also connected to posterior retinotopic areas identified by population receptive field mapping^64^ conducted in the same participants (Extended Data Fig. 9c,d; Extended Data Fig. 10), which presumably provide low-level visual inputs to FFA. The relatively widespread “off-diagonal” connections of the FFA sites are in contrast with some NHP studies reporting connectivity largely confined to the face patches^62,63^, but are consistent with the central role of face processing in human cognition, including the need to integrate visual social information with linguistic information as well as the need to make complex social inferences based on visual social signals^65–67^.

Our findings also inform the connectivity of the brain’s language regions^68^. Extended Data Fig. 12 shows two language-dominated stimulation sites (green crosses), both of which showed strong connections with other language-processing regions (Extended Data Fig. 12d,j). Site 7-3 in the left LPFC was also connected to a speech perception fROI in the superior temporal gyrus (Speech.STG, dark brown) and the visual word form area (Word.VWFA, dark green) in the remote ventral temporal cortex. It was also connected to several MD fROIs in nearby lateral and medial prefrontal cortices, which showed mixed MD and speech responses. Site 9-1 in the left anterior temporal cortex was connected to face (Face.fSTS; f=fundus) and TOM (TOM.TPJ) fROIs in the dorsal temporal cortex, both of which showed strong language responses alongside their fROI-defining functions, as well as to an MD fROI in the lateral temporal cortex; the latter connectivity may be better explained by its strong episodic projection responses shared with the site.

Extended Data Fig. 13 shows two MD-dominated sites^4,69^, both of which showed strong connections with other MD regions (Extended Data Fig. 13d,j; also see highlighted MD fROIs in Extended Data Fig. 13). Site 1-3 in the right LPFC was additionally connected to an object perception fROI in ventral temporal cortex (Obj.VTC), which showed relatively low MD responses. The site was also connected to a nearby language fROI (Lang.IFG). Site 8-2 in the right dorsal prefrontal cortex connected to a lateral temporal object perception fROI (Obj.LTC), and two physical reasoning fROIs (Phys.PC and Phys.FC) that both showed strong MD responses.

To synthesize the observations across all stimulation sites, we constructed an fROI-site connectivity matrix and an fROI-to-fROI connectivity diagram (Fig. 5). We first assigned each stimulation site to the fROI that best overlapped with its nearby voxels (Fig. 5a). This matrix is sparse: most sites covered only one or two fROIs, justifying the unique assignment. Compared to the contrast-level activation matrix (Fig. 3a), this sparsity reflects the fact that fROIs were defined for only a subset of the cortical locations activated by a given function. Consequently, at locations where multiple functions overlap, we may not have fROIs nominally associated with all of them. Relatedly, an fROI nominally associated with one function may also contain voxels activated by other functions, as illustrated by the example sites (Fig. 4 and Extended Data Figs. 11-13). The mixed nature of responses within fROIs is an important feature of cortical organization that should be taken into account when interpreting fROI-level results.

**Figure 5.**
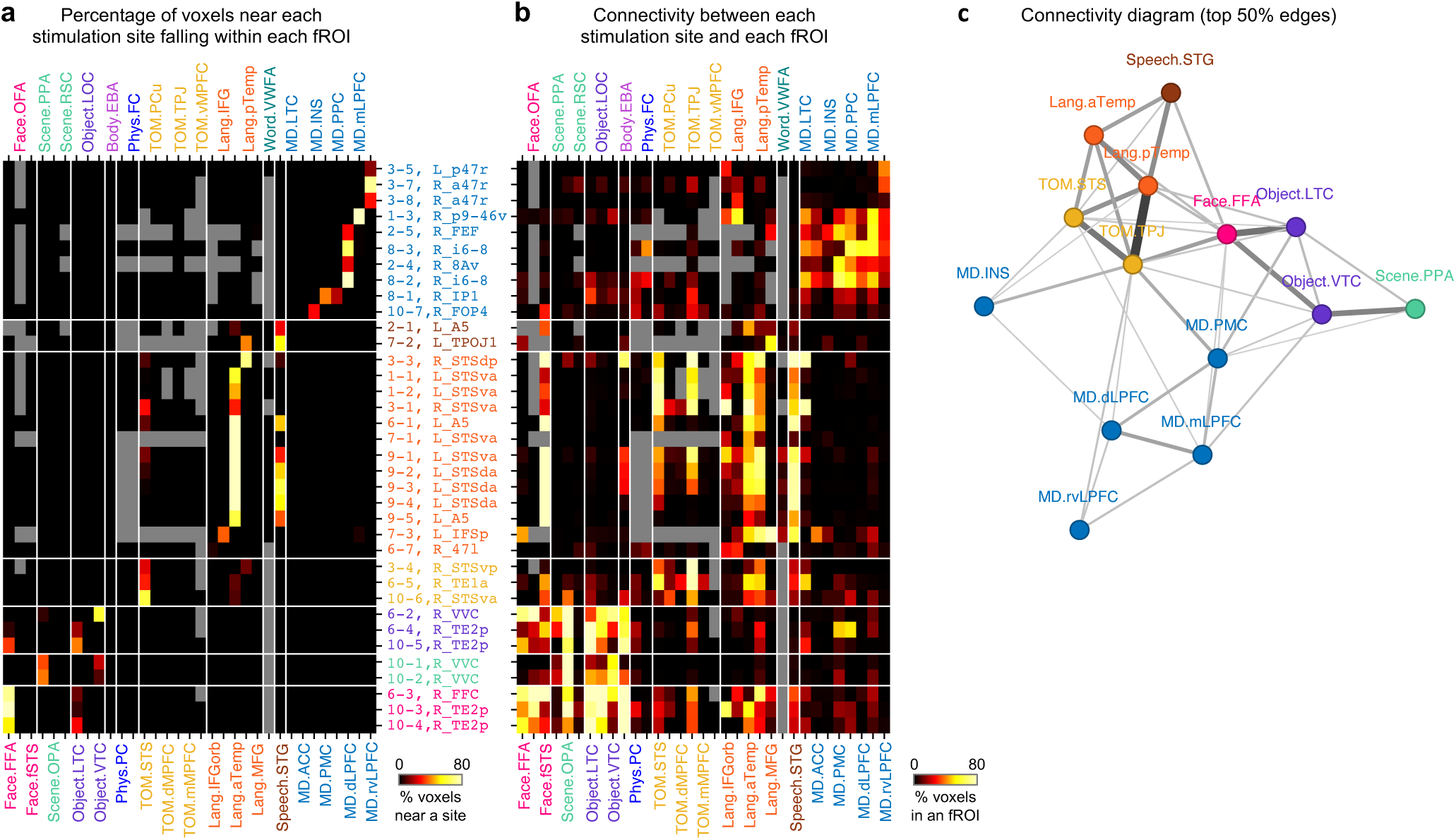
fROI-level connectivity. (a) fROI×site matrix showing the percentage of voxels near each stimulation site falling within each fROI. Gray entries indicate missing data, i.e., when an fROI was undefined, such as the right VWFA, or wasn’t identified in a subject. (b) fROI×site matrix showing the percentage of connected voxels within each fROI for each stimulation site. (c) fROI-to-fROI connectivity diagram between a subset of fROIs in (a,b). Lines connect fROI pairs with above-threshold (50th percentile) connectivity; line thickness indicates connection strength.

Figure 5b shows the connectivity of each stimulation site to each fROI (i.e. the percentage of connected voxels within each fROI), revealing several notable patterns that echo the individual examples (also see Extended Data Fig. 14 for results merged across stimulation sites of the same fROI). First, among the visual processing fROIs, the FFA was connected to the other two face fROIs (OFA and fSTS), to the EBA for body perception^51^, to multiple object fROIs, and to OPA, a posterior scene fROI^70^, but not to the nearby scene-selective PPA fROI^50^. In addition, the FFA was connected to the TPJ TOM fROI (and, to a lesser extent, the STS TOM fROI) and to several language fROIs (mainly the aTemp and pTemp fROIs in the temporal lobe, which are closer to FFA than other language fROIs, e.g., the prefrontal ones, consistent with the distance dependence of connectivity). Second, the PPA fROI was also connected to the Scene.OPA fROI and multiple object fROIs, but, in contrast to the FFA, showed much weaker connectivity to face, TOM, and language fROIs. Third, the language fROIs were strongly connected to each other and, additionally, to the STG speech fROI. They also showed abundant connections with the fSTS face fROI and two TOM fROIs (TPJ and STS), which often showed some degree of language response (as site 9-1 in Extended Data Fig. 12), but they had very sparse connections with MD fROIs. Fourth, the MD fROIs were strongly connected to each other and also modestly, yet diversely, to fROIs supporting non-MD functions, in line with the idea that the MD network needs to integrate information from multiple systems in the service of goal-directed behaviors^4^.

To derive an fROI-to-fROI connectivity matrix (see METHODS), for each pair of fROIs, we identified all entries in Fig. 5b of which rows (with each site assigned to an fROI) and columns corresponded to that pair, averaged them, and forced the resulting matrix to be symmetric. We took the top 50% strongest connections in the matrix to derive an fROI-to-fROI connectivity diagram (see METHODS). The resulting diagram (Fig. 5c) contains a subset of fROIs due to the still-incomplete coverage of our data, yet it nonetheless recapitulates the main findings from Fig. 5b, most notably the tight interconnections between fROIs that support face processing, TOM, and language functions.

### Compare rsFC and es-fMRI connectivity

We next examined how anatomical connectivity relates to rsFC patterns. Because site-seeded rsFC is a widely used metric of connectivity in human neuroscience^12,35,53,59^, we first compared it (computed from the same participants’ individual resting-state data) with the anatomical connectivity derived by es-fMRI at the same stimulation sites. We found only a modest association.

Figures 6a and 6b show examples of rsFC seeded with two stimulation sites (6-5 and 6-6) in the same participant’s right hemisphere. Both sites appeared to reside in the default-mode network (DMN) based on canonical rsFC patterns that were highly similar between the two sites, despite their being located far apart. In contrast, their connectivity patterns derived by es-fMRI (Fig. 6c,d) were dissimilar: most of each site’s connections were localized in different subsets of locations within the high-rsFC regions, with some connections also corresponding to low rsFC. Another example pair of sites (6-3 and 6-4; Extended Data Fig. 15), adjacent to each other on the ventral temporal surface, showed similar rsFC patterns but distinct connectivity patterns, neither of which was well captured by rsFC.

**Figure 6.**
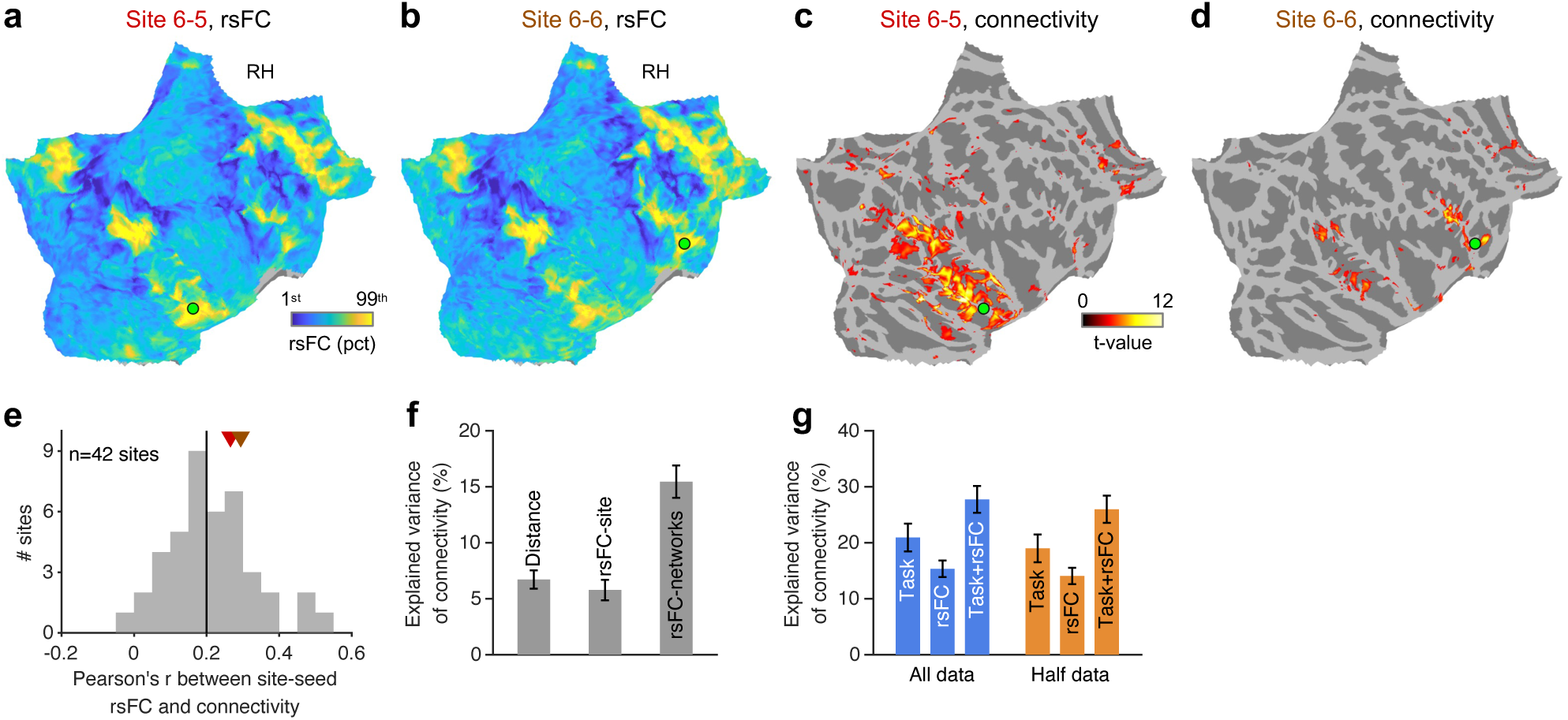
RsFC versus anatomical connectivity. (a,b) RsFC seeded with two stimulation sites (6-5 and 6-6) in the same participant (S6), displayed on flattened cortical surfaces. (c,d) Es-fMRI connectivity patterns for the same two sites. (e) Histogram of the Pearson correlation between site-seeded rsFC and es-fMRI connectivity (excluding voxels near the site) across all 42 stimulation sites. Vertical line indicates the median. Colored triangles mark the example sites. (f) Cross-validated variance explained by distance, site-seeded rsFC, and network-seeded rsFC. (g) Cross-validated variance explained by task-based, rsFC (network-seeded), and combined models using full and half (odd runs only) data. All error bars denote SEM across sites.

Figure 6e summarizes the correlation (Pearson’s r) between site-seeded rsFC and es-fMRI connectivity across all 42 stimulation sites, which was only moderate (median/mean/std=0.20/0.22/0.12). Both example sites in Fig. 6a-d were close to the median correlation. Site-seeded rsFC predicted connectivity slightly worse than distance (Fig. 6f; rsFC-site, 5.8% explained variance vs. Distance, 6.7%), consistent with the moderate correlation.

Previous studies have suggested that multivariate rsFC features can predict task-based functional maps^71–75^. Since the task-based functional profile showed a reasonably strong association with anatomical connectivity (Fig. 3c), we asked whether multivariate rsFC features could serve as a task-free functional profile for predicting connectivity. We derived these features by calculating rsFC between each voxel and 12 individually defined intrinsic functional networks (Extended Data Figs. 16 and 17; see METHODS) described in the Cole-Anticevic Brain-wide Network Partition^76^. In contrast to site-seeded rsFC, which measures a single correlation between each voxel’s time course and the stimulation site’s time course, network-seeded rsFC characterizes each voxel by a vector of 12 correlations with major functional network time courses, capturing a voxel’s position within the brain’s large-scale functional architecture. Network-seeded rsFC predicted connectivity substantially better (Fig. 6f; rsFC-networks, 15.5%) than site-seeded rsFC (5.8%).

Combining task-based activations and network-seeded rsFC produced a moderate increase in performance (Fig. 6g, All data; Task/rsFC/Task+rsFC = 20.9%/15.3%/27.7%; using 37 functionally defined sites), indicating that resting-state fMRI carries non-overlapping predictive power for connectivity relative to task fMRI, and vice versa. This increase was not attributable to the added scan time by using data from both modalities, because a combined model using features derived from half runs (odd runs only) of both modalities (Fig. 6g, Half data; Task/rsFC/Task+rsFC = 19.0%/14.1%/26.0%) still outperformed models using full-length data from either modality alone.

## DISCUSSION

By combining es-fMRI with comprehensive task-based and resting-state fMRI in the same individuals, we mapped close-to-ground-truth anatomical connectivity in the human cortex and related it to its functional organization. The central finding is that connectivity obeys an asymmetric functional-similarity principle: connected regions are likely to share a functional profile, but, because connectivity is sparse, regions with similar profiles are unlikely to share a connection. Mapping connections across individually defined functional regions revealed converging patterns of information flow through well-studied regions for face processing, language, theory of mind, and other cognitive functions.

An important implication concerns functional integration across cognitive systems. Because task fMRI activations often show some degree of spatial overlap across functions, connections associated with shared activations with the stimulation site may simultaneously link the stimulation site with regions associated with functions beyond those shared with the site. The fusiform face area (FFA), for instance, was connected not only to other classic face-responsive regions but also to the theory of mind region in the temporoparietal junction and to the anterior temporal language areas, and connected voxels in the latter two regions exhibited some response to faces alongside responses to their preferred functions. Thus, connections associated with shared functions and the mixed selectivity of cortical regions may together support integration across functional systems. At the same time, a non-trivial number of connected locations showed no shared activations with the stimulation sites, indicating that anatomical connectivity can support integration across functional systems in the absence of a shared function.

The “off-diagonal” connections we found between regions that support distinct functions, such as connections between areas activated most strongly by face processing and areas activated most strongly by theory of mind reasoning, begin to lay the foundation for understanding cross-network information transfer in the human brain. The FFA showed connections with a variety of other functional regions, including areas activated by object processing (posterior to the FFA) and TOM areas in the TPJ and STS. The former connections could supply the FFA with visual input (e.g., object features) important for face recognition, in contrast to earlier arguments that the face processing pathway can function normally when the object recognition pathway is severely compromised^77^, while the latter connections could be important for using face information in downstream social cognition. Future work could follow this “visual features to faces to social information processing” pathway^65,78,79^. The moderate connections of the FFA with some speech and language areas fit well with both the role of lip-reading during speech decoding^80,81^ and the role of facial expression in interpreting language (e.g., sarcasm and humor)^82^.

Another unexpected finding was the set of connections between TOM regions in the TPJ and STS and the temporal language areas. Although the language and TOM networks have been shown to have some degree of functional correlation, especially pronounced during language comprehension^83^, these connections could also reflect the verbal nature of the TOM contrasts and the possibility of linguistic confounds^84^. This result needs further exploration in future studies, including with non-verbal TOM paradigms^85^.

The multiple demand^4^, or executive function, fROIs (including MD.dlPFC, MD.mLPFC, MD.rvLPFC, MD.PMC, and MD.INS) showed strong connections mainly with each other, despite being widely distributed across the brain, which could underlie their importance for a very wide variety of cognitive tasks. Notably, two of these MD regions (MD.mLPFC, MD.PMC) also showed moderate connections with areas activated in object processing tasks (Object.LTC, Object.OTC), paralleling anatomical findings in primates^6,86^. These two MD areas may be the primary recipients of information important for interacting with objects, as well as the source of feedback connections important for top-down control of visual processing.

Why do anatomical connections tend to target regions with similar functional profiles? One possibility is that the sharing of information via anatomical connections across different cortical areas leads to functional convergence, through experience-dependent plasticity. For example, anatomical connections between a site activated in TOM tasks in the TPJ and a site activated in TOM tasks in the STS might have promoted even closer functional properties in the two areas. Co-activation in common tasks could lead to plasticity within cortical areas akin to Hebbian learning^87^. Another possibility is that genetic or developmental programs may cause sites with shared functional properties to target one another anatomically. In this case, the “learning’ may take place primarily over evolutionary timescales. The two possibilities are not mutually exclusive: developmental wiring may establish a coarse functional-similarity bias that experience-dependent plasticity then refines.

Resting-state functional connectivity, despite its name, proved to be a poor proxy for anatomical connectivity at the level of individual stimulation sites, warranting interpretive caution in seed-based analyses. However, characterizing voxels by their rsFC profiles across multiple intrinsic networks substantially improved prediction for es-fMRI connections, and combining these features with task data yielded further predictive gains. This finding suggests that metrics capturing a region’s functional profile, whether derived from tasks or resting-state network affiliation, can predict aspects of its anatomical connections.

Several limitations should be noted. Es-fMRI lacks laminar resolution, and the directionality of connections cannot be established, although prior work suggests that es-fMRI PBRs correspond to forward (orthodromic) transmission^42^. Our coverage, while spanning many functional systems, remains incomplete; broader, denser sampling will be needed to construct a comprehensive connectivity map of the human brain and to test the principle of topographic mappings as we have done in our NHP work^6^. Our participants are epilepsy patients, and although their task-based activations were broadly consistent with those of neurotypical adults, we cannot fully rule out effects of epilepsy-related reorganization. Additionally, although we defined functions using well-established localizers, individual topographic variability and confounds inherent in some task contrasts could affect functional assignments and complicate interpretation. The population-level findings are robust to these issues because neither functional similarity nor linear regression analyses were sensitive to the purity of individual task contrasts, whereas the fROI-level connectivity diagram, which is still preliminary, should be interpreted with caution. As we examine specific functional systems in greater detail in future work, careful validation of localizer accuracy will be essential.

Functional connectomics^88^, which measures how functionally defined neural elements are wired together, is crucial for understanding the brain’s computations. In animals, connectivity can be probed at neuronal or synaptic resolution, including in recent multi-modal connectomes^89,90^. In humans, the millimeter-scale resolution we achieve here, while far coarser, is well-suited to reveal the pathways of information flow across functional regions and to link connectivity with functional organization. Absent a synaptic-level connectome paired with function in humans, the mesoscale connectivity achievable with es-fMRI, paired with functional profiling, offers the best route to building more accurate and complete models of human cognition^91^.

Looking ahead, es-fMRI can be used to test many circuit-level predictions about how functional regions communicate, moving beyond correlational measures toward causal characterization of information flow. Excitingly, this approach enables the development of complementary directions in clinical neuroscience, as patient-specific es-fMRI maps could inform intracranial neuromodulation strategies^92–95^.

## METHODS

### Participants

Eight participants with medically refractory epilepsy undergoing invasive diagnostic investigation by means of weeks-long stereoelectroencephalography (SEEG) monitoring at Massachusetts General Hospital (MGH) were enrolled. They also completed noninvasive MRI sessions at MIT when they were not implanted (before, after SEEG monitoring, or both). All procedures were approved by IRBs at MGH and MIT, and all participants provided written informed consent. Electrode placement was based solely on clinical considerations. The participants showed high cooperation during the MRI sessions, and their task-based fMRI results were comparable to those of neurotypical adults (Extended Data Fig. 4).

### Safety testing

Before conducting es-fMRI in any participant, we performed extensive phantom-based heating tests to assess RF-induced heating of implanted metal components. We filled phantoms with polyacrylic acid (PAA) gel mixed with NaCl, which has physical, thermal, and electrical properties similar to those of human brain tissue^96^, and measured temperature at SEEG electrode contacts and bone screws using fiber-optic probes (OSENSA PRB-500). We identified the distance between extension cables and the RF transmit coil as a key, underappreciated factor: with cables positioned at the scanner isocenter using a custom-made track, heating remained negligible (<0.5 °C) for T1-weighted and EPI sequences across three 3T scanners (Siemens Prisma, Skyra, and Vida). High-SAR sequences (T2-weighted) caused greater heating, and poor cable placement (e.g., when the cable lies on the floor of the MRI bore or is taped to a CP transmit-receive coil) could lead to severe heating (>10 °C within 2 minutes). Based on these results, we used only low-SAR sequences with isocenter cable positioning for all patient scans.

### Electrical stimulation

A subset of SEEG electrodes (PMT Corporation, MN, USA) for epilepsy monitoring was used for stimulation. Most electrodes were removed at the bedside prior to scanning, retaining 2-4 electrodes containing stimulation sites. Stimulation was delivered using constant-current bipolar stimulation between adjacent contacts, using charge-balanced biphasic pulses (200 us cathodal phase, 100 us blank, 200 us anodal phase). The amplitude was set at 8 mA in the first two participants, and then switched to 7.5 mA to ensure the charge density was below the manufacturer’s recommended upper limit (30 μC/cm^2^). To verify that stimulation was safe and did not induce discomfort, we tested all stimulation sites at the bedside before es-fMRI scans. In two stimulation sites (8-2 and 8-3), we reduced the amplitude to 4 mA to eliminate stimulation-related sensations. We used a block design with alternating 16-second null (no stimulation) and stimulation blocks (272 sec per run; nine/eight rest/stimulation blocks). We collected one run per stimulation site, except for sites 6-3, 8-1, 8-2, 8-3, for which we collected two runs per site. In each stimulation block, a ∼210 ms pulse train containing 70 pulses (one pulse every 3 ms, or 333 Hz) was delivered once per second. Participants were instructed to close their eyes during es-fMRI scanning. A licensed physician was present to monitor for after-discharges and side effects for the duration of the study. At the amplitude level used for es-fMRI, participants reported no sensations during bedside stimulation and es-fMRI.

### MRI data acquisition

The es-fMRI sessions were conducted using a Siemens 3T Vida scanner with a 64-channel coil at MGH. T1-weighted structural images (3D MPRAGE, 1 mm isotropic) and single-echo gradient-echo EPI functional images (TR=0.8 s, TE=33 ms, flip angle=52°, 2.5 mm isotropic, 64 slices, multiband factor=8) were acquired. Field maps were also acquired as a pair of spin-echo EPI images with reversed phase-encode directions for TOPUP distortion correction.

The noninvasive MRI sessions were conducted using a Siemens 3T Prisma scanner with a 32-channel coil at MIT. Task and resting-state functional images were acquired using a single-echo gradient-echo EPI sequence (TR=1.0 s, TE=35.2 ms, flip angle=60°, 2.0 mm isotropic, 72 slices, multiband factor=6). The localizer battery included 13 tasks producing 32 contrasts targeting 13 functions (Extended Data Table 1). T1-weighted structural images (3D MPRAGE, 0.8 mm isotropic) and field maps (paired spin-echo EPI images) were also acquired.

### fMRI data analysis

All functional data were preprocessed using fmriprep^97^ 24.1.1. Processing steps included motion correction and fieldmap-based distortion correction. Spatial smoothing was performed in the T1w space (individual volume; FWHM=3 mm).

General linear models were fitted using FS-FAST (selxavg3-sess) with the SPM canonical hemodynamic response function (-spmhrf 0). Task fMRI GLMs included no temporal filtering to avoid regressing out low-frequency components in the task design of some localizers; es-fMRI GLMs included a high-pass filter at 0.008 Hz (the simple AB-blocked design was not susceptible); resting-state fMRI GLMs included a high-pass filter at 0.008 Hz and a low-pass filter at 0.1 Hz. For all GLMs, we included rigid-body motion parameters as nuisance regressors, framewise-displacement-based timepoint censoring, and polynomial detrending (-polyfit 1 for es-fMRI and task fMRI to avoid regressing out task design, -polyfit 2 for resting-state fMRI).

Statistical maps were projected from T1w to fsnative (individual surface) using FreeSurfer^98^ mri_vol2surf (--projfrac-avg 0 1 0.2, trilinear interpolation) and then to fsaverage (template surface) using FreeSurfer mri_surf2surf with no further smoothing. We refer to the surface vertices as voxels in the paper.

Statistical maps were thresholded at p<0.001 uncorrected for es-fMRI and p<0.01 uncorrected for task fMRI. The minimum absolute t-value corresponding to this threshold in T1w volume was used to threshold the projected t-value on the surface. The residual time courses of resting-state fMRI GLMs were used to compute resting-state functional connectivity using Pearson’s correlation.

### Electrode localization

The location of electrode contacts was manually confirmed by co-registering post-implantation CT images with pre-operative clinical structural MRI images. The clinical structural MRI images were then aligned to the T1w space using rigid transformations. The mid-point of contact dipoles was projected to nearest mid-thickness vertex of individual surface and then transformed to fsaverage surface. When planning stimulation sites, we chose dipoles with mid-points and both contacts in the gray matter or the white matter adjacent to gray matter. We did not find excessively stronger/more es-fMRI activations when the stimulation sites were in white matter compared to when they were in gray matter, unlike what we observed in NHPs^6^. We defined nearby voxels (≤4 mm) based on the Euclidean distance between 3D coordinates on inflated surfaces.

### Es-fMRI connectivity metric

The connectivity metric was the es-fMRI t-statistic for the stimulation-versus-null contrast. Voxels with p>=0.001 were set to zero; above-threshold t-values (both positive and negative) were preserved. We focused on positive BOLD responses (PBRs) as connections, and set negative values that passed the threshold to zero in most cases (except in Extended Data Fig. 1c). Voxels with positive t-values (after thresholding) were deemed “connected”, and those with zero t-values were deemed “unconnected”.

### Task fMRI activation metric

The task fMRI activation metric was the task fMRI t-statistic for each functional contrast of interest. Voxels with p>=0.01 were set to zero; above-threshold t-values (positive only) were preserved, corresponding to “activated” voxels.

### Effective SNR

To compare statistical power across studies, species, and fMRI modalities, we normalized t-values to the same scan duration. With everything else held constant, t-values are linearly proportional to the square root of scan duration. To ensure a fair comparison across datasets with different total scan times, we defined effective SNR as t_raw_/sqrt(actualScanTime/272), where 272 seconds (the run length of es-fMRI) serves as the normalization unit. We then took the 99th percentile of positive t-values (significant before normalization; p<0.0001 to match the threshold used for NHP data) for each stimulation site or task contrast. The comparison was conducted in individual volume space for all datasets, which have isotropic resolutions of 2-2.5 mm. For task fMRI, we only considered between-condition contrast in two-condition localizers (e.g., emotional pain > physical pain in TOM pain variant), which matches es-fMRI.

### Criteria for functionally defined sites

Of the 42 stimulation sites, 37 were classified as functionally defined based on having at least 20% of nearby voxels (≤4 mm) activated in at least one contrast (78.3±25.7% in these sites). The remaining 5 sites were excluded from analyses requiring task-based functional profiles but included in analyses that did not.

### Functional similarity metric

We computed the cosine similarity between the functional profile of a stimulation site and each voxel. Before computing, we merged contrasts belonging to the same function: for each function group, we computed the fraction of contrasts in the group with positive activation at each voxel (value 0-1), not considering unavailable contrasts (Extended Data Table 2), rather than averaging raw t-values. The 32 contrasts were mapped to 13 functions (see Extended Data Table 1): Face (3 contrasts), Body (2), DMN (3), TOM (2), Episodic memory (1), Language (3), Speech (1), Articulation (1), MD (4), Intuitive physics (1), Scene (2), ObjA (4), ObjB (5). Voxels not activated in any contrast were labeled as unknown.

### On-surface distance

Geodesic distance between cortical locations was approximated as great-circle arc length on the FreeSurfer registration sphere (?h.sphere, radius=100 mm), rescaled so that the sphere surface area (both hemispheres combined) matches the white-matter surface area.

### Linear regression analysis

We used ordinary least squares regression (MATLAB’s fitlm) to predict voxel-wise es-fMRI t-values from various predictor sets. Predictors were not standardized prior to fitting. To fairly compare models with different numbers of predictors, we performed leave-one-area-out cross-validation using the HCP MMP1.0 parcellation^99^. For each stimulation site, the model was trained on voxels of all ipsilateral HCP areas except one and tested on voxels of the held-out area, iterating over all areas. Voxels near the site were excluded from training and testing. Model performance was quantified as 1 minus the normalized mean squared error (NMSE) of predictions on held-out data, computed by concatenating predictions from all held-out areas into a single vector per site. Unique and partial variance contributions were computed by comparing nested models: unique variance for predictor X equals R^2^_full model_ minus R^2^_model without X_; partial variance for X equals R^2^ of a model containing only X.

### Functional regions of interest (fROIs)

fROIs were defined as the intersection of population-based anatomical masks and individual-based task fMRI activations for the corresponding function. For most fROIs, we used anatomical masks curated by the efficient multifunction fMRI localizer (EMFL) study^58^, including face, scene, and body ROIs^57^, language ROIs^5^, TOM ROIs^100^, word ROI^101^, and speech and physics ROIs. For MD fROIs, masks from Fedorenko lab^102^ were usually used, sometimes merging adjacent ones into larger regions, and in some cases supplemented with or purely based on HCP areas according to the core and extended MD network definition^69^: INS (insula per Fedorenko lab’s naming), PMC – posteromedial cortex (postparietal), PPC – posterior parietal cortex (midparietal, antparietal), dLPFC – dorsolateral prefrontal cortex (supfrontal), mLPFC – mid LPFC (precentral_a_precg, midfrontal, precentral_b_ifgop), rvLPFC – rostroventral LPFC (midfrontalorb; HCP areas a9-46v, p10p, a10p, p47r, a47r, 11l), LTC – lateral temporal cortex (HCP areas TE1p, TE1m), and ACC – anterior cingulate cortex (HCP areas SCEF, 8BM, a32pr, d32). No parcels were included to define DMN or episodic memory fROIs.

In-mask voxels showing activation in any of the contrasts of the defining function of an fROI were considered part of that fROI. We excluded fROIs with <50 voxels to reject spurious results. For face, scene, body, and word fROIs, a voxel was included if it showed activation in any fROI-defining contrasts AND did not show activation in any inverted fROI-defining contrasts. For example, voxel included in a scene fROI should be activated in Eff.S-O or DyObj.S-O (or both) AND not activated in either Eff.O-S or DyObj.O-S. Conversely, for object fROI, a voxel was included if it showed activation in any fROI-defining contrasts (ObjA and ObjB) AND did not show activation in any defining contrasts for face, scene, body, or word fROIs. N-back object-perception contrasts (Face.NBack, ObjB-F.NBack) were not used in the fROI definition due to potential confounding by attention level.

### fROI-to-fROI connectivity diagram

To derive the fROI-to-fROI connectivity diagram, we first computed the weighted average of connectivity (Extended Data Fig. 14) between stimulation sites of one fROI (rows in Fig. 5b) and all fROIs (columns in Fig. 5b). A stimulation site was assigned to the fROI with the highest percentage coverage of the site’s nearby voxels (Fig. 5a). We forced the resulting fROI-to-fROI connectivity matrix to be symmetric by averaging the asymmetric matrix and its transpose. Pairwise connectivity values were then available for all fROIs containing stimulation sites plus one remaining fROI, for which we chose TOM.TPJ for its functional importance. The top 50% of non-zero connectivity values (50th percentile threshold) across the selected subset of fROIs were retained for the diagram (Fig. 5c). Node positions in the diagram were determined by a force-directed embedding based on connectivity values. Line width is proportional to connectivity strength.

### Individualized rsFC network parcellation

Starting from the population-based Cole-Anticevic Brain-wide Network Partition^76^, which assigns cortical areas to 12 large-scale rsFC networks, we iteratively adjusted voxel-wise network assignments to match individual rsFC patterns, following a prior study^103^. At each iteration, we computed the mean resting-state time course for each network seed, reassigned each voxel to the network with which it had the highest Pearson correlation, and updated the seed time courses using the new network assignments. A spatial cluster of fewer than 40 same-network voxels is merged into the largest neighbor network. This procedure was repeated for up to 20 iterations on the fsaverage6 surface, or until convergence (fewer than 1% of voxels changed their network assignment compared to the previous iteration).

### Data availability

The data that support the findings of this study are available from the corresponding authors upon request.

## ACKNOWLEDGMENTS

We thank all patient-participants for their time and effort in completing this study. We also thank Karen A. Rich, Rebecca A. Larosa, Christopher Whirty, Kara J Houghton, Fausto Minidio, Grace Ng, Jacqueline Campos, and Bradford Sheldon for their assistance. A.B., E.F., and R.M.R. were partially supported by NINDS U01 NS121471. N.G.K. was partially supported by NIMH UM1 MH130981-01.

## AUTHOR CONTRIBUTIONS

Conceptualization: R.X., R.D., and R.M.R. Formal Analysis: R.X., E.F., N.G.K., and R.D. Investigation: R.X., A.B., A.T., N.S., A.W., S.H., A.K., C.N., N.J., A.M., A.F., S.D., E.H., C.C., and R.M.R. Methodology:

R.X., A.B., A.T., N.S., A.W., S.H., V.K., C.V., J.K., and R.M.R. Resources: R.X., A.T., N.S., A.K., P.N., S.Y., J.K., and R.M.R. Supervision: R.X., A.B., R.D., and R.M.R. Writing – original draft: R.X. and R.D. Writing – review & editing: R.X., A.B., V.K., E.F., N.G.K., R.D., and R.M.R.

---

**Extended Data Fig. 1.**
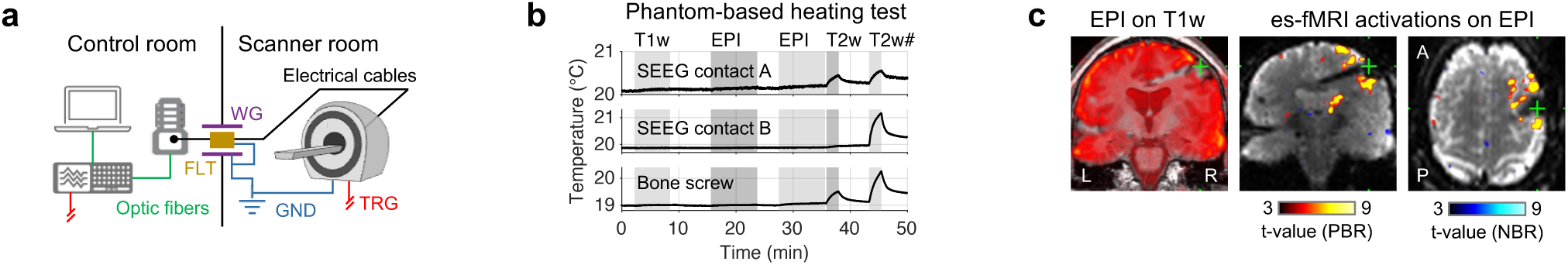
Es-fMRI setup and safety validation. (a) Schematic of the es-fMRI setup. Isolated electrical stimulation was delivered from the control room via extension cables connected to implanted SEEG electrodes with participants staying in the scanner room. In the control room, a computer interfaces with a Tucker-Davis RZ5D processor, which in turn interfaces with a Tucker-Davis Subject Interface stimulator, both via optic fibers. A waveguide (WG) provides access into the scanner room. An RF-filter (FLT) sharing ground with the scanner and waveguide reduces external RF noise introduced to the scanner room. A trigger signal (TRG) from the scanner synchronizes stimulation with MRI acquisition. (b) Phantom-based heating test. Temperature measured by fiber-optic sensors at two SEEG electrode contacts and a bone screw (top to bottom panels) over time during sequential MRI acquisitions (T1w, EPI, T2w; T2w# corresponds to the same T2w sequence, but with an extension cable layout different from the standard es-fMRI setup). With cables positioned at the scanner isocenter using a custom-made track (standard es-fMRI setup), heating remained negligible (<0.5 °C) for T1w and EPI sequences. (c) Example es-fMRI data. Left: EPI image overlaid on T1w anatomy, showing good functional-to-anatomical alignment. Mid/right: example coronal/axial slice of es-fMRI activations overlaid on the EPI volume, showing both PBRs and NBRs (absolute t-value≥3).

**Extended Data Fig. 2.**
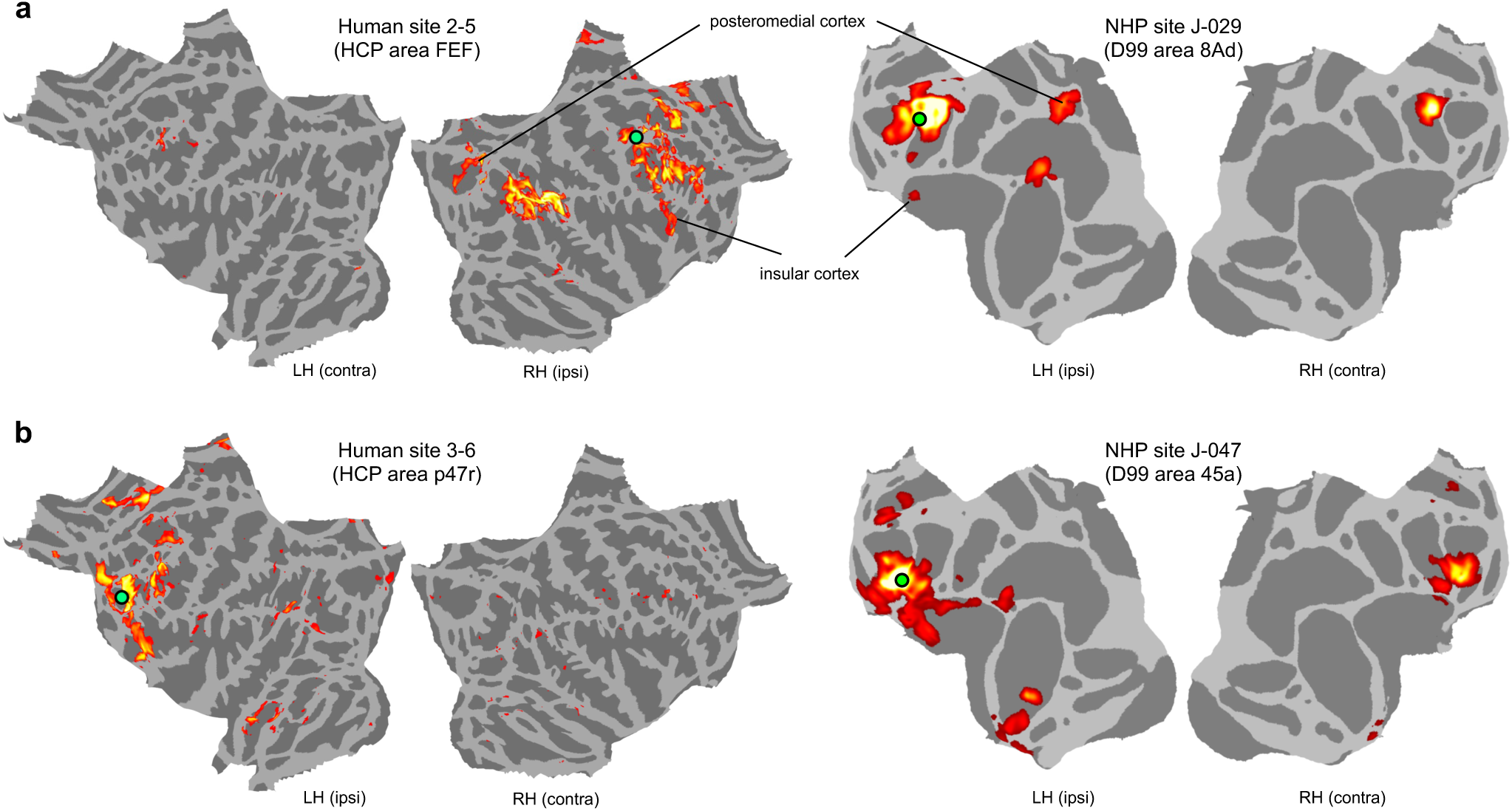
Cross-species comparison of es-fMRI connectivity patterns. Two additional pairs of stimulation sites (see Fig. 1d for a third pair) at matching cortical locations in humans and NHPs. Each pair shows PBRs on flattened surfaces for the human site and the corresponding NHP site.

**Extended Data Fig. 3.**
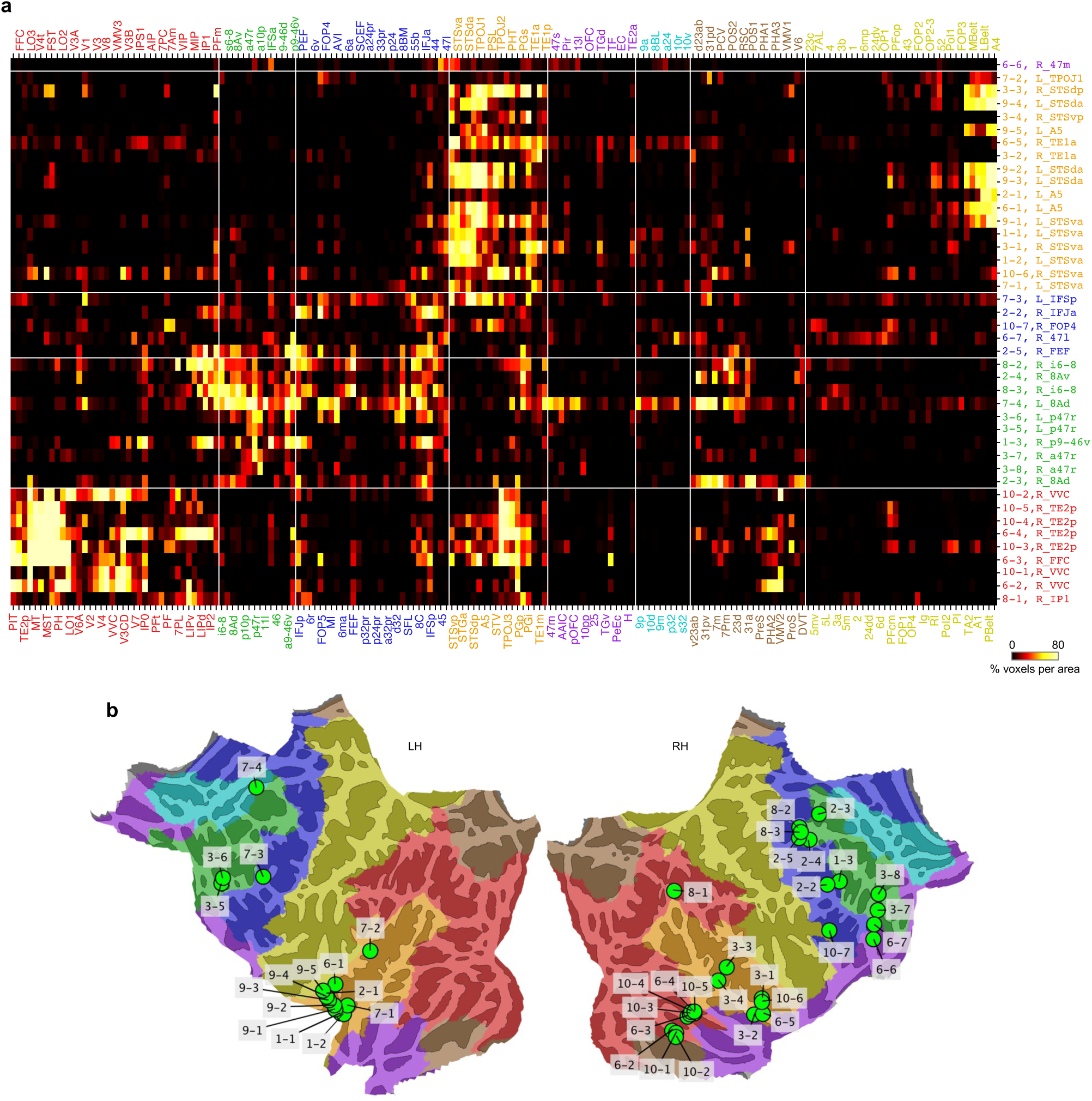
Area×site connectivity matrix and stimulation site locations. (a) Connectivity matrix between ipsilateral HCP areas (columns) and all 42 stimulation sites (rows), displayed as the percentage of voxels showing PBRs in each of the 180 ipsilateral HCP areas. Sites and areas are grouped by major cortical parcels and sorted by spatial adjacency within each parcel. The parcels are derived by merging adjacent HCP areas with similar task and resting-state fMRI responses. (b) Locations of all stimulation sites on the flattened cortical surface. Text colors in (a) match overlay color of parcels in (b).

**Extended Data Fig. 4.**
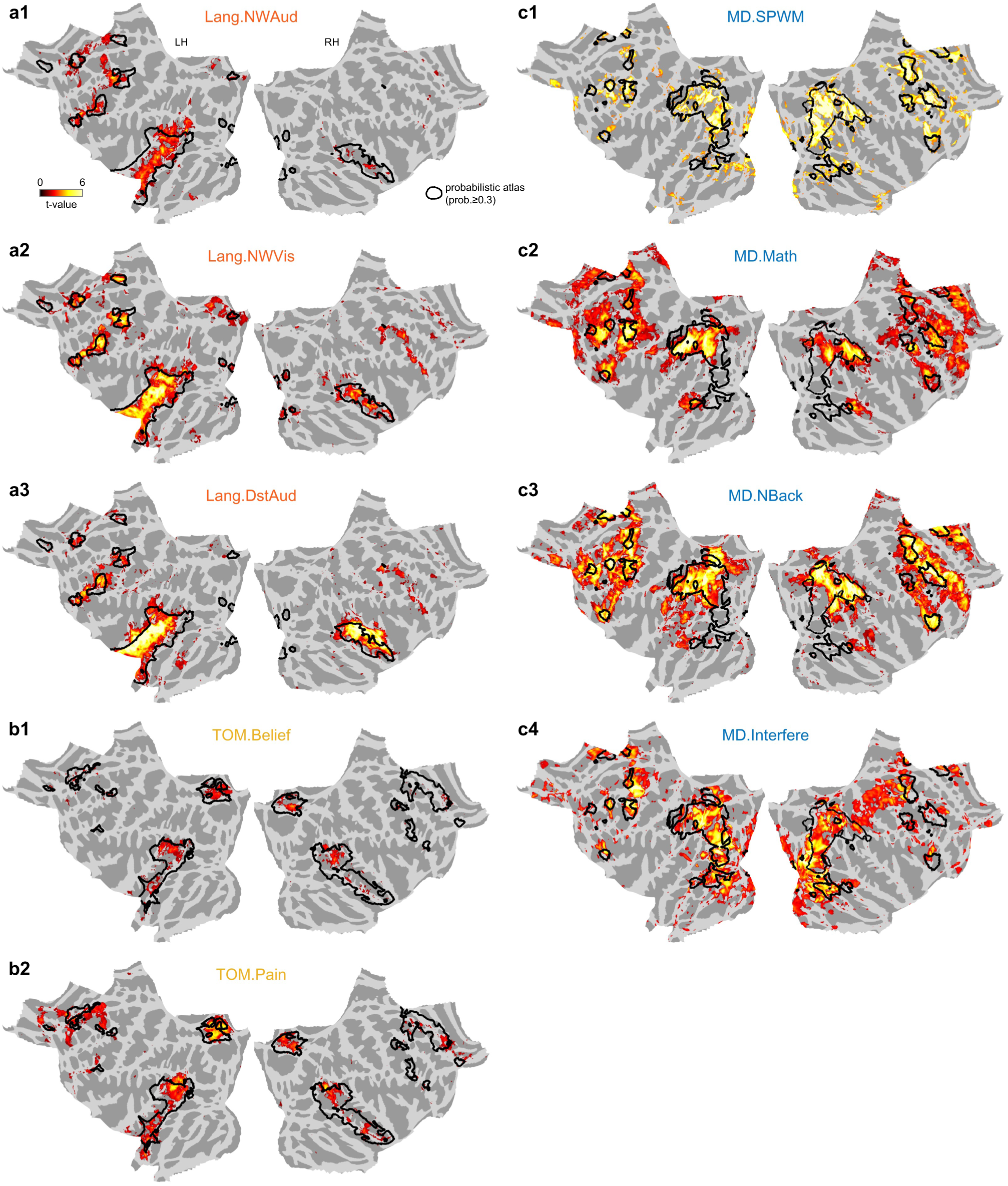
Validation of task-based fMRI localizers. Subject-averaged activation patterns (mean t-value≥1) from individual language, TOM, and MD contrasts in the present participant cohort are shown alongside published probabilistic atlases derived from neurotypical adults^108^. Strong activations in our participants were well co-localized with high-probability zones in the population atlases (black contours). Activation patterns of contrasts for other functions were also broadly consistent with expectations from the literature.

**Extended Data Fig. 5.**
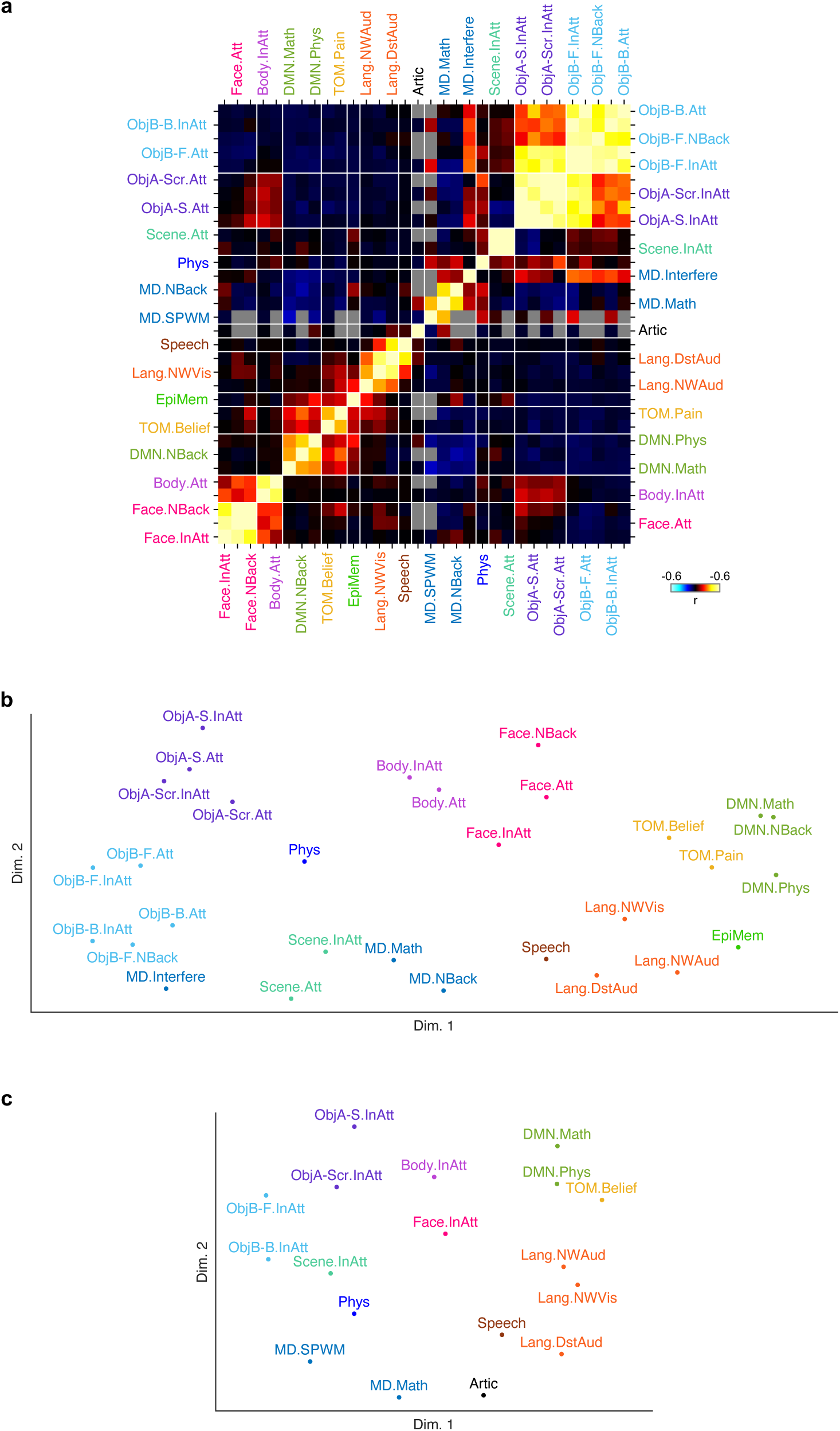
Cross-contrast correlation of task fMRI activation patterns. (a) Correlation matrix showing pairwise Pearson correlations between cortex-wide activation patterns across all 32 task fMRI contrasts. Contrasts are ordered by function as in Fig. 3a. As expected, contrasts targeting the same function show highly correlated activation patterns, while cross-function correlations are generally much lower. (b,c) Two-dimensional visualization of inter-contrast distances, computed separately for two maximal cliques of contrasts. A clique is a maximal subset of contrasts in which every pair has defined correlation across subjects (i.e., at least one participant had both contrasts of the pair available). Clique 1 (b) contains the larger subset; clique 2 (c) contains a second, partially overlapping subset. Within each clique, coordinates were obtained by running UMAP (n_neighbors=15, min_dist=0.3, n_epochs=5000, spectral initialization) for 100 rounds on the mean correlation distance matrix, computing pairwise Euclidean distances from each UMAP embedding, taking the element-wise median across rounds, and applying non-metric MDS (stress criterion, 100 random restarts) to the median distance matrix to produce the final 2D layout. In both panels, contrasts are clustered by function, i.e., contrasts targeting the same function are positioned nearby.

**Extended Data Fig. 6.**
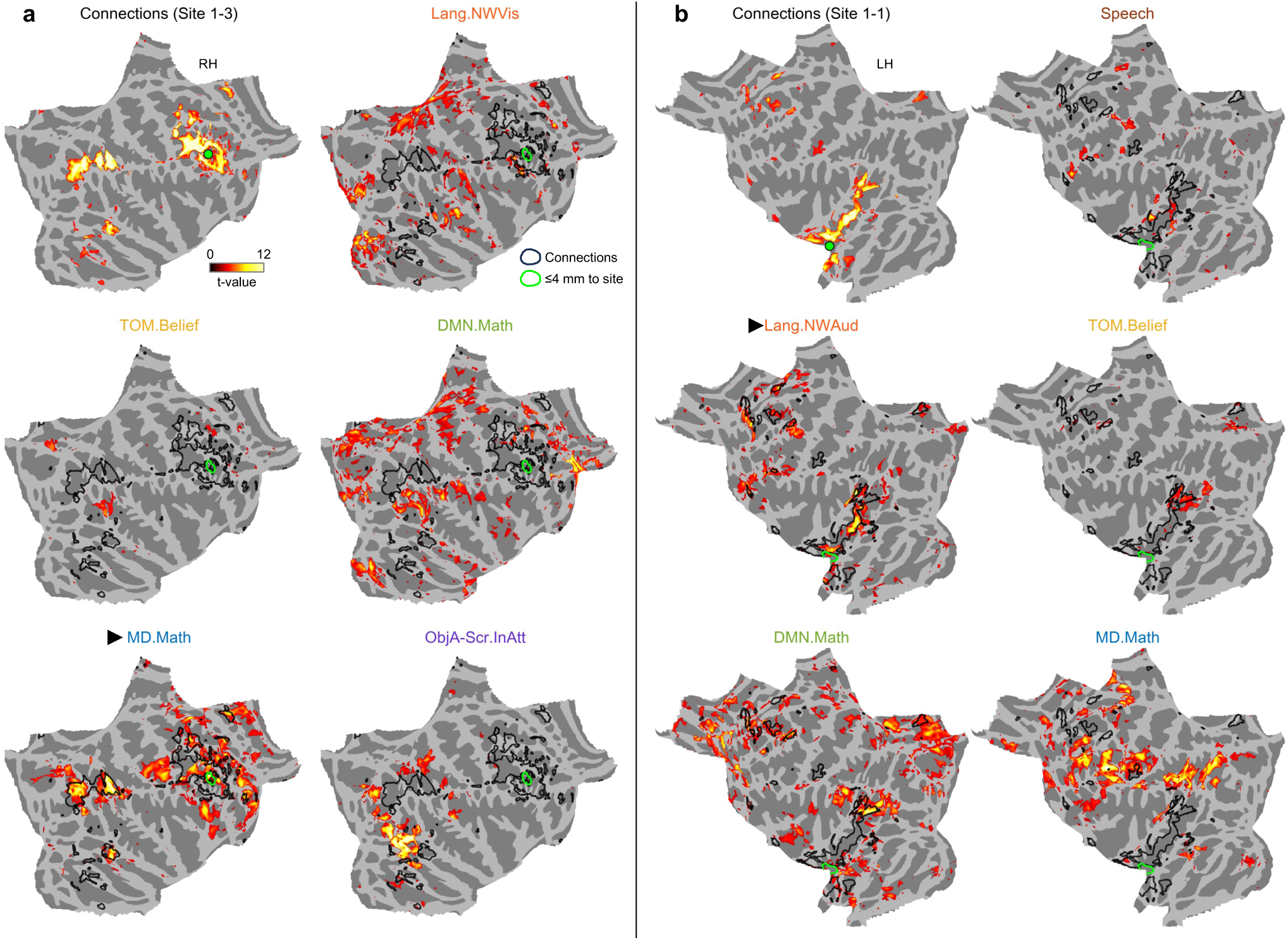
Example stimulation sites illustrating the correspondence between connectivity and function. (a) Stimulation site 1-3 (subject S1, site 3). Top-left: es-fMRI connections displayed on flattened cortical surface. Other panels: task fMRI activation maps for selected contrasts in Language, TOM, DMN, MD, and ObjA functions, with black contours indicating es-fMRI connections. The site was activated in an MD contrast (MD.Math), and most connected regions also show MD activations, while largely avoiding activations in non-MD contrasts. (b) Stimulation site 1-1 (subject S1, site 1). The site was predominantly activated in a language contrast (Lang.NWAud), and its connections overlap best with activations of the same contrast. Both examples illustrate the common observation that connectivity tracks function.

**Extended Data Fig. 7.**
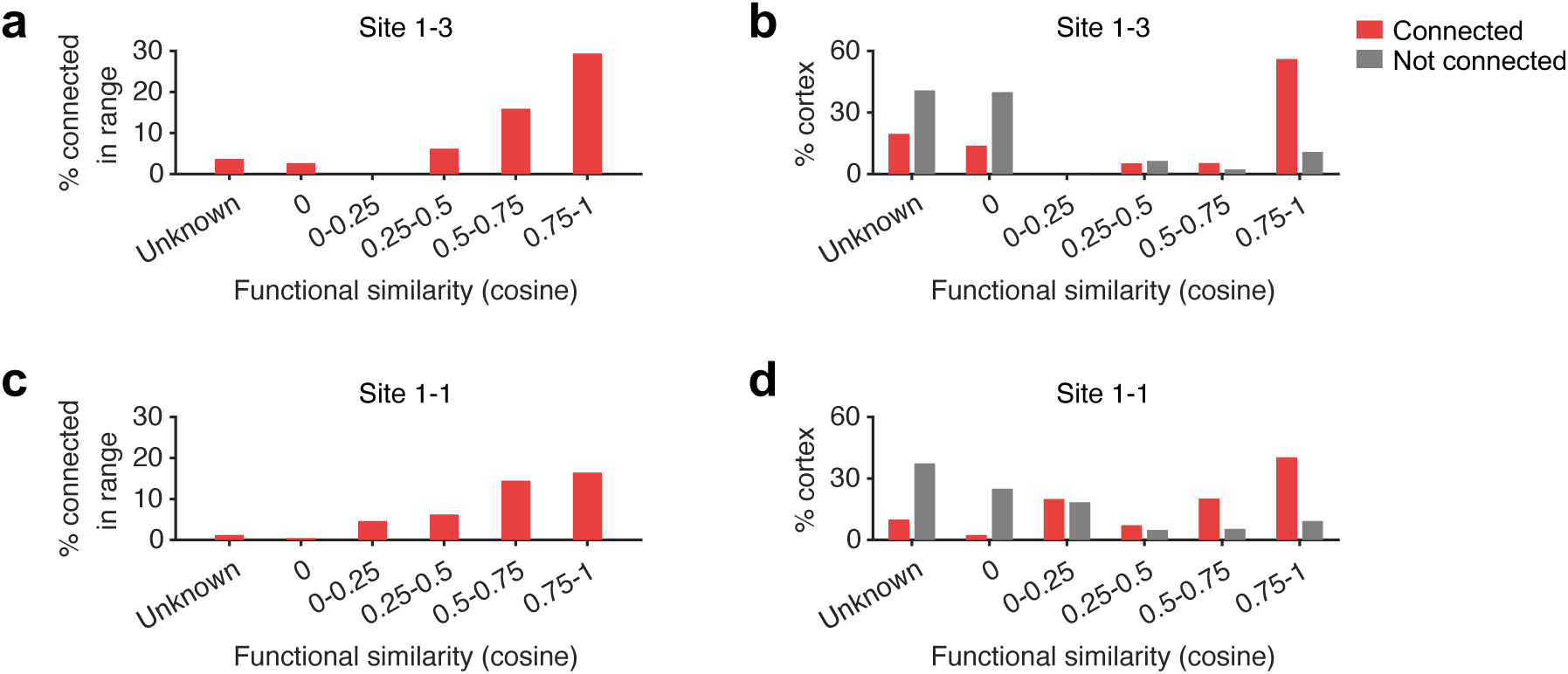
Functional similarity statistics for two example stimulation sites. (a,c) Percentage of connected voxels across functional similarity ranges at sites 1-3 (a) and 1-1 (c). (b,d) Distribution of functional similarity for connected (red) and unconnected (gray) voxels at sites 1-3 (b) and 1-1 (d). The results are broadly consistent with population-average results (Fig. 2f,h), with variations in exact numbers.

**Extended Data Fig. 8.**
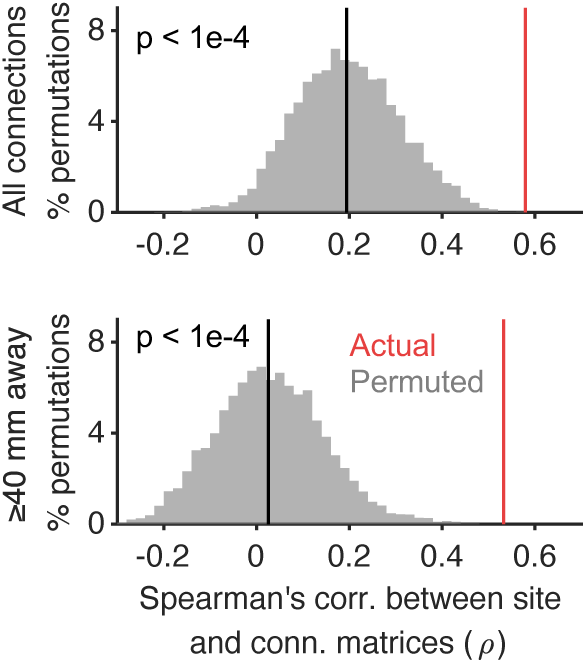
SPIN permutation tests of the connectivity-function correspondence. Histogram of SPIN-permuted (20000 permutations) similarity values (Spearman’s ρ) between two contrast×site matrices in Fig. 3a,b. Top: using all cortical voxels, excluding those near the site. Bottom: excluding voxels <40 mm from stimulation sites. Red lines indicate the actual value. Black lines indicate the median of SPIN-permuted values.

**Extended Data Fig. 9.**
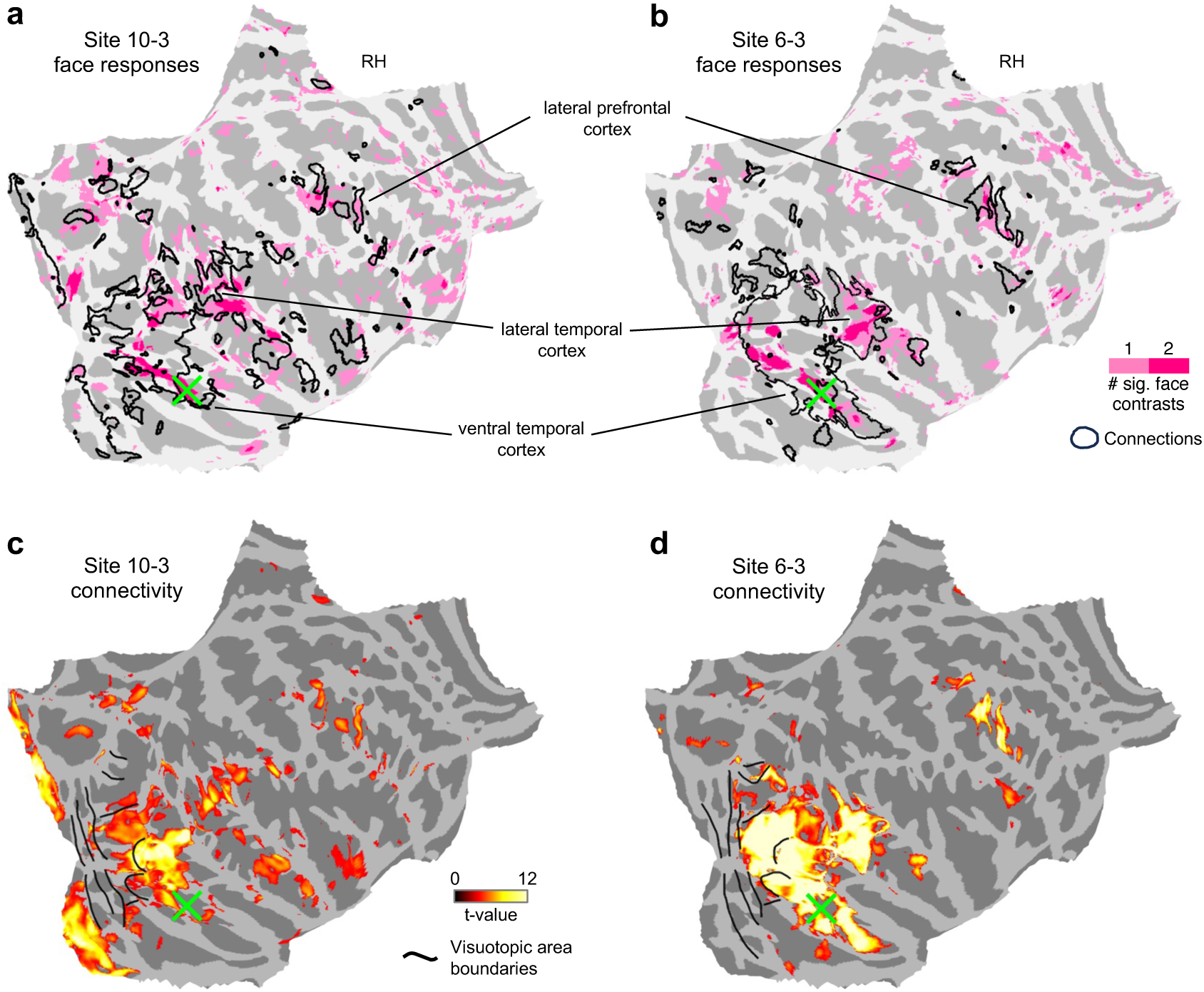
Connectivity of two FFA stimulation sites in comparison to face activations and retinotopic areas. (a,b) Connectivity patterns (black contours) of FFA sites 10-3 (a) and 6-3 (b) (green X) with face-contrast task fMRI activations (colored overlay) on flattened cortical surfaces. Both sites connect to face-responsive regions in the ventral temporal, lateral temporal, and lateral prefrontal cortices. (c, d) Connectivity of FFA sites 10-3 (c) and 6-3 (d) as colored overlay with retinotopic area boundaries in black contours, derived from the same-individual population receptive field mapping results. For both FFA sites, connected retinotopic areas included LO1, LO2, and regions near the MT complex (Extended Data Fig. 10).

**Extended Data Fig. 10.**
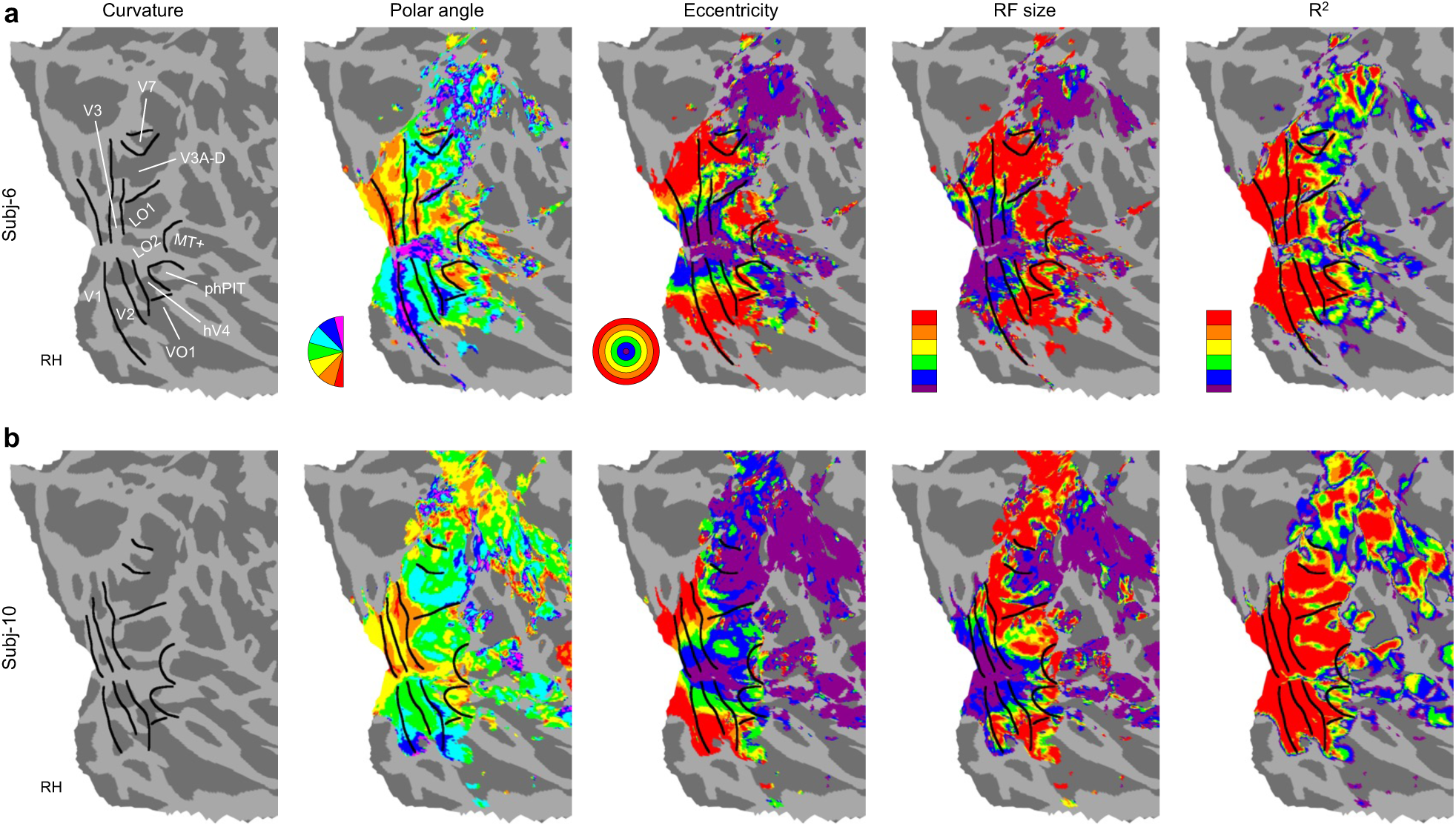
Population receptive field (pRF) mapping. Retinotopic maps for the two participants with FFA stimulation sites (S6 and S10). Voxel-wise results of polar angle, eccentricity, RF size, and goodness of fit of pRF models (R^2^) are displayed on flattened cortical surfaces, with area boundaries drawn as black lines. These maps were used to tentatively identify retinotopic areas, including V1, V2, V3, LO1, LO2, MT complex (MT+), hV4, VO1, phPIT, V3A-D, and V7 regions, following criteria and naming conventions in previous studies^109,110^. The pRF analysis was performed using the analyzePRF toolbox^111^. The max diameter of pRF stimuli (from the website of analyzePRF) was ∼17.5° for S6 and ∼13.7° for S10, and the viewing distance was 137 cm for both participants.

**Extended Data Fig. 11.**
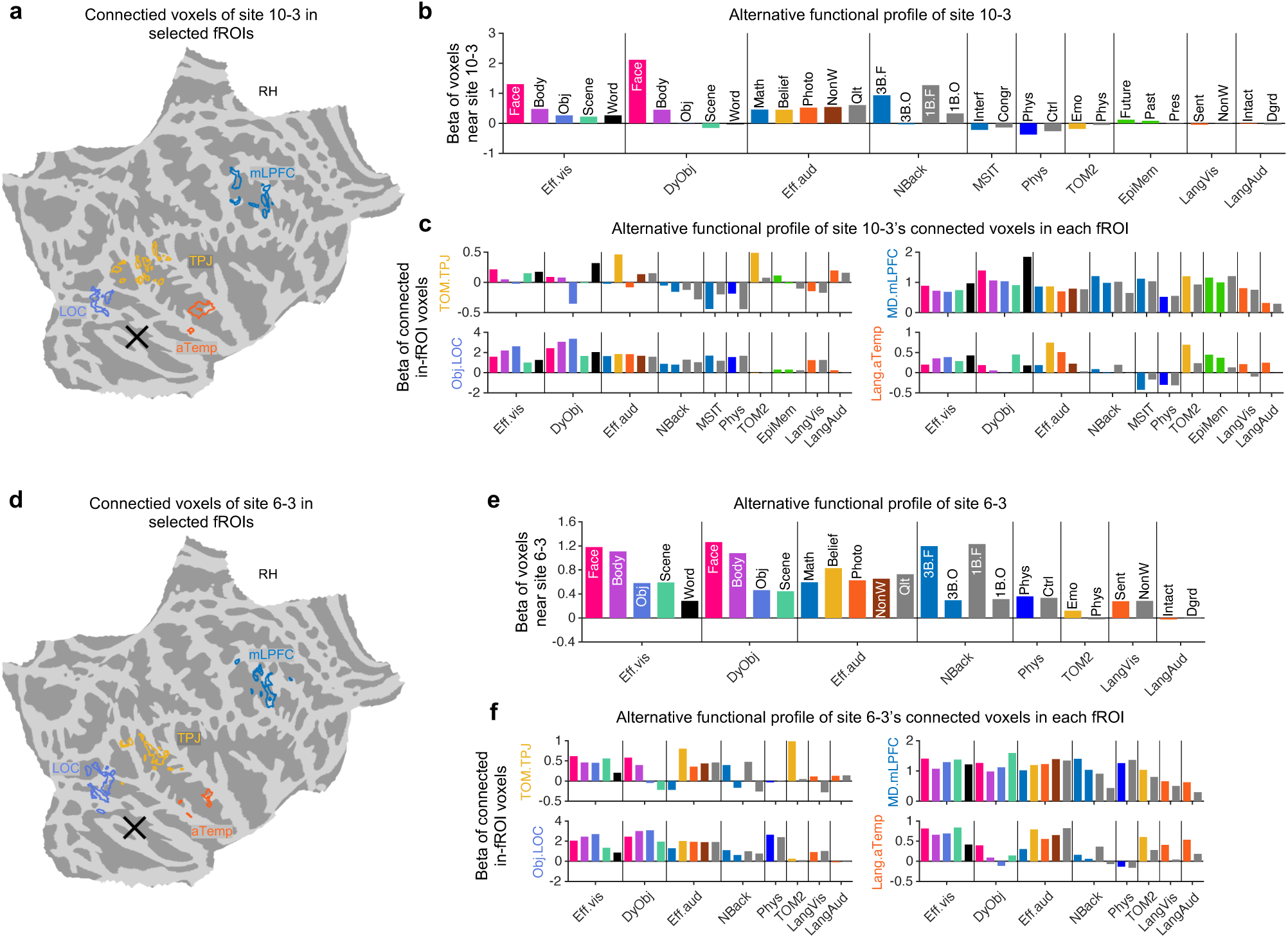
GLM-derived Beta weights of two FFA stimulation sites and connected fROIs. Functional characterization of two FFA stimulation sites (10-3 and 6-3) and their connected fROIs, using condition-level beta weights rather than the contrast-level metric shown in Fig. 4. (a) Contours of connected voxels of site 10-3 within fROIs on the flattened cortical surface, with black X indicating the stimulation site’s location. Colors of contours match fROI names in (c). (b) Alternative functional profile of site 10-3: mean beta weights of all voxels near the site per condition. Conditions are sorted by localizer. For condition interpretations, refer to contrast definitions in Extended Data Table 1. Conditions are colored based on the pertinent function. (c) Alternative functional profile of connected voxels of site 10-3 within each of the four fROIs: mean beta weights of all connected voxels per condition. Conditions are in the same order as (b). (d-f) Same as (a-c) for site 6-3.

**Extended Data Fig. 12.**
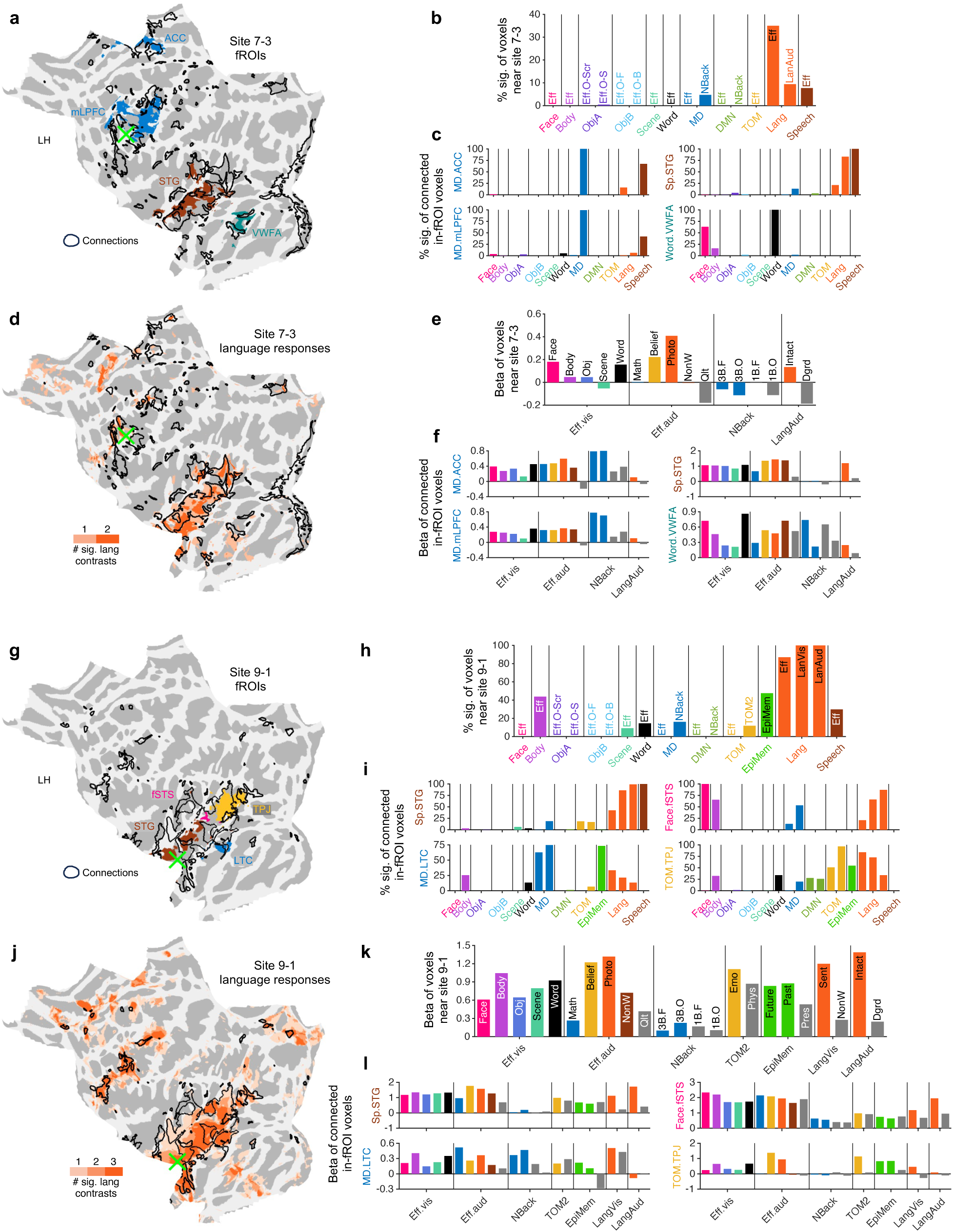
Connectivity of two language-dominated stimulation sites. (a) Stimulation site 7-3 in the left LPFC (green X) shown on the flattened cortical surface with fROIs as colored overlays and connections as black contours. (b) Functional profile of site 7-3 as assessed by the percentage of voxels near the site activated in each contrast. (c) Functional profile of connected voxels within each of the four fROIs as assessed by percentage of connected voxels activated per contrast. (d) Connectivity patterns (black contours) of site 7-3 (green X) with language-contrast task fMRI activations (colored overlay). (e) Alternative functional profile of site 7-3 as assessed by mean beta weights of all voxels near the site per condition. Conditions are grouped by localizer (Extended Data Table 1). (f) Alternative functional profile of connected voxels within each of the four fROIs as assessed by mean beta weights of all connected voxels per condition. (g-l) as (a-f), results of stimulation site 9-1 in the left anterior superior temporal sulcus.

**Extended Data Fig. 13.**
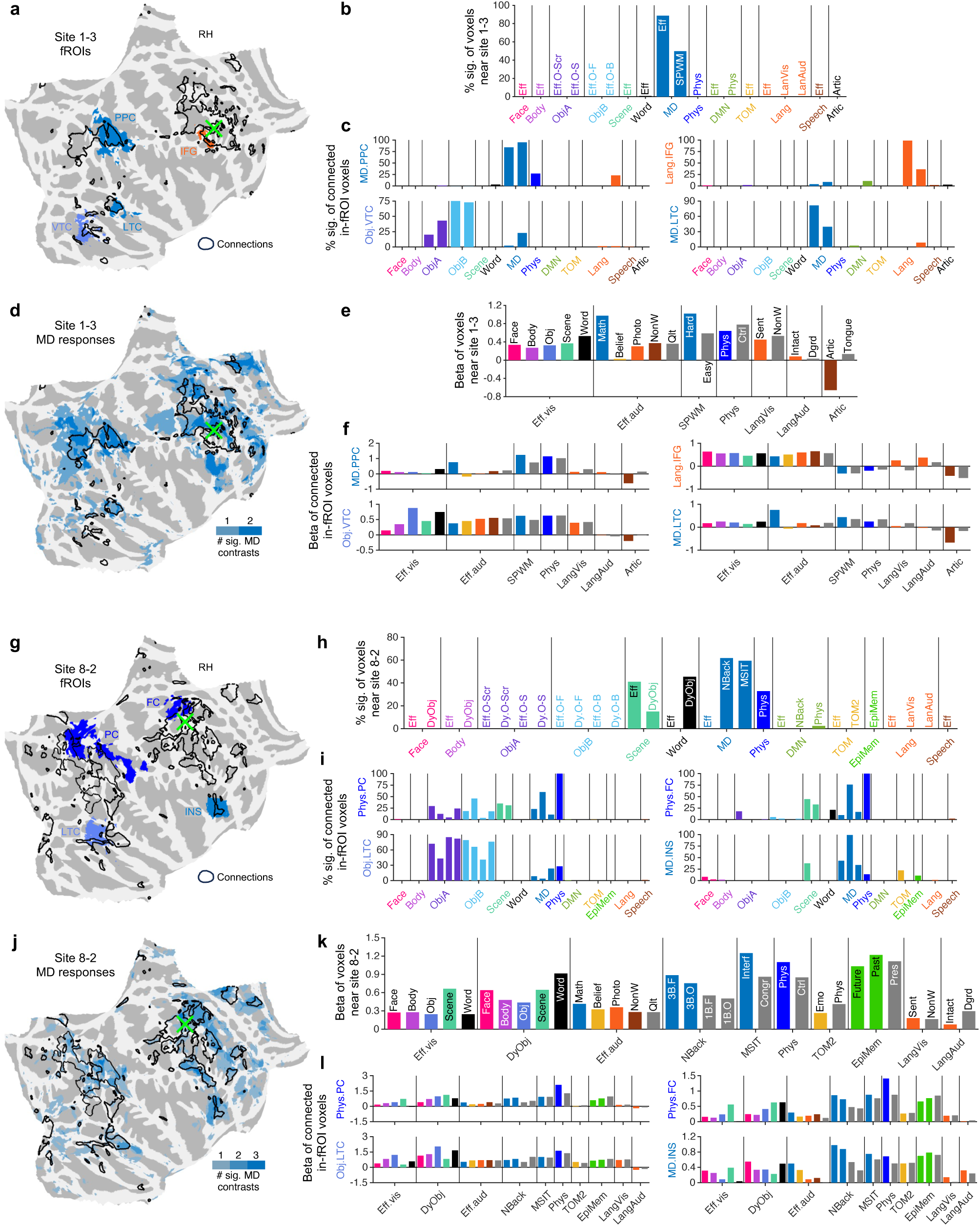
Connectivity of two MD-dominated stimulation sites. Same format as Extended Data Fig. 12 for two MD-dominated sites, 1-3 (a-f) and 8-2 (g-l).

**Extended Data Fig. 14.**
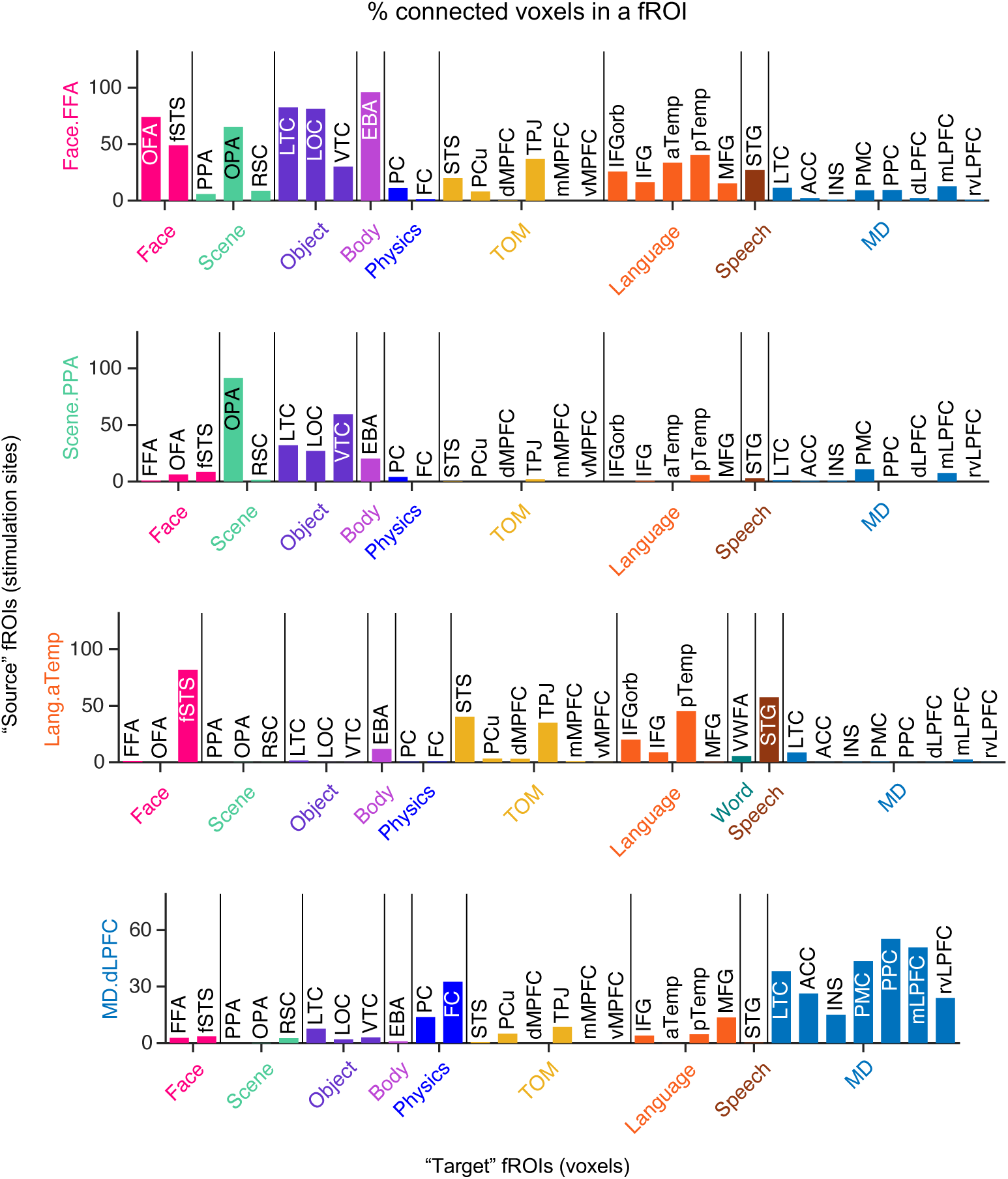
Rows of fROI-level connectivity matrix merged across stimulation sites of the same fROIs. Each panel shows the connectivity from one source fROI to all target fROIs as the percentage of voxels in each target fROI that are connected to the (source) stimulation sites. The percentage was weighted-average across sites assigned to the source fROI (weights proportional to each site’s coverage of the source fROI, i.e., entries in Fig. 5a). Top to bottom: Face.FFA, Scene.PPA, Lang.aTemp, and MD.dLPFC as the source fROI. Target fROIs are grouped by function. Bars corresponding to missing data are omitted.

**Extended Data Fig. 15.**
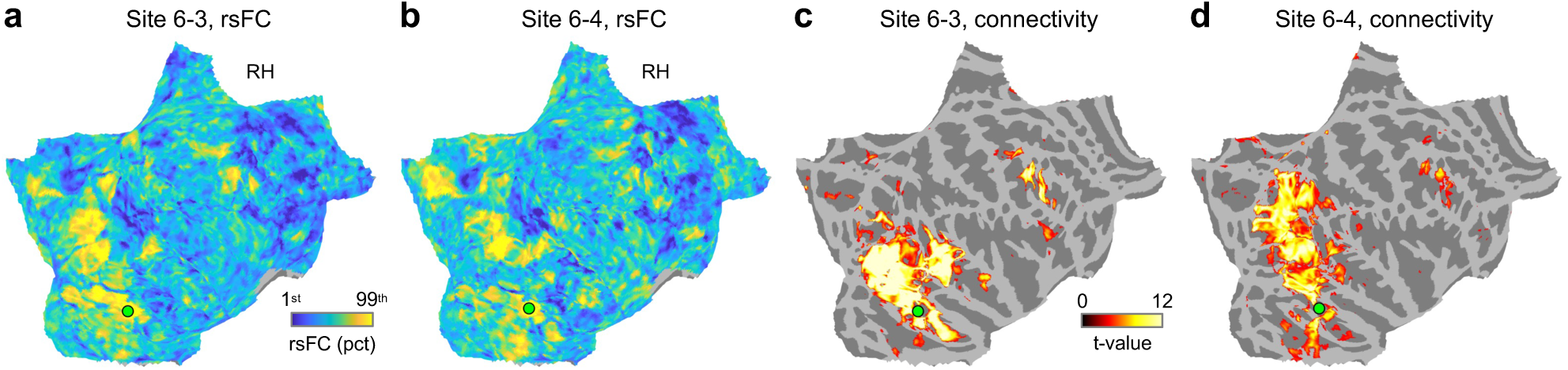
Comparison between rsFC and anatomical connectivity at two additional example sites. (a,b) RsFC seeded with two stimulation sites (6-3 and 6-4) in the same participant (S6), displayed on flattened cortical surfaces. (c,d) Es-fMRI connectivity patterns for the same two sites.

**Extended Data Fig. 16.**
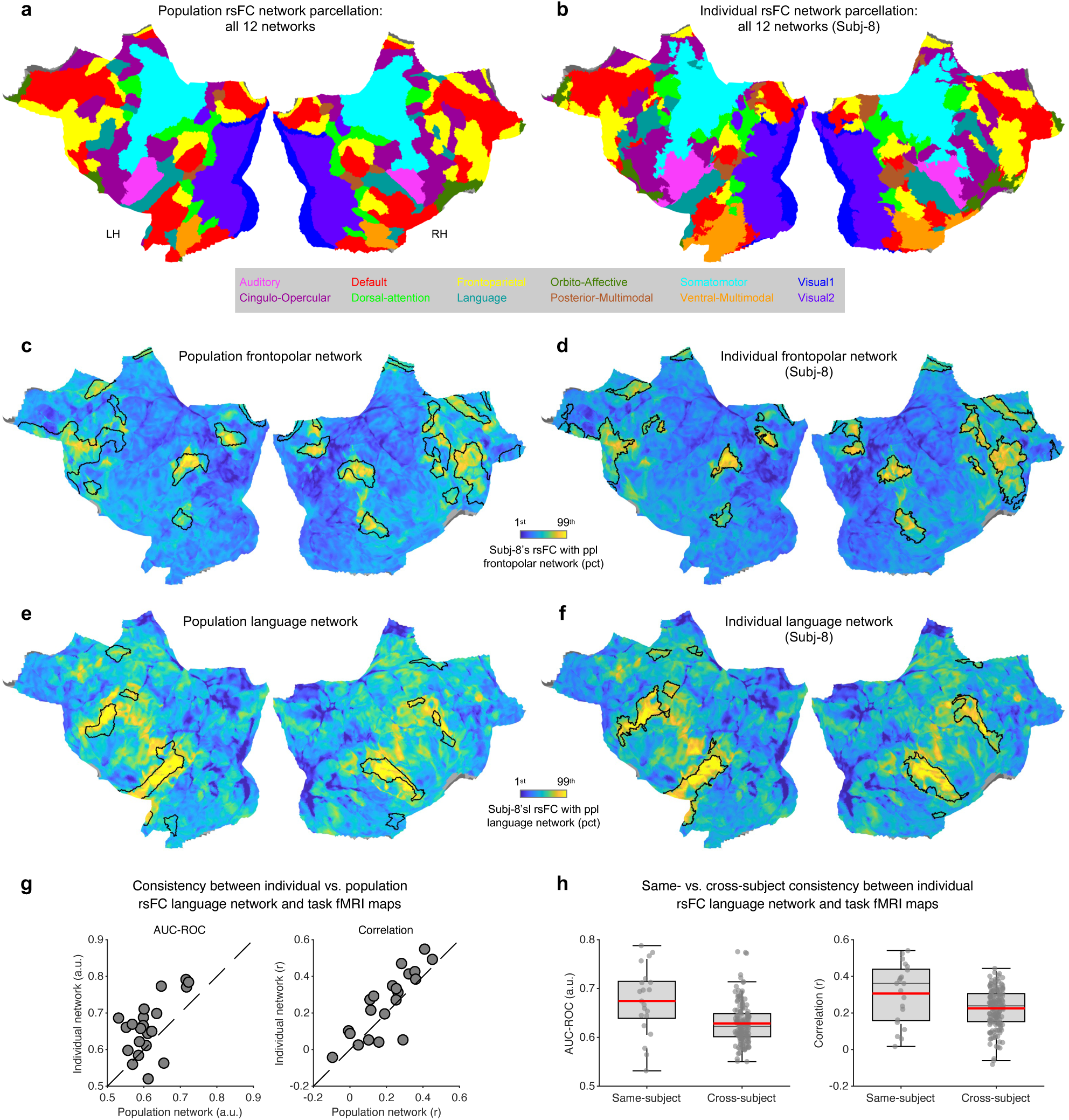
Individualized rsFC network parcellations. (a) Population-based Cole-Anticevic Brain-wide Network Partition (CAB-NP; 12 networks). (b) RsFC network parcellation derived from resting-state fMRI data of example participant S8, starting from the CAB-NP parcellation. (c,d) RsFC with the population frontopolar network, with boundaries of the population (c) and individualized (d) frontopolar networks shown as black contours. The individualized network boundaries aligned better with the rsFC pattern, despite the latter being seeded with the population network. (e,f) As (c,d), for the language network. (g,h) To validate the individualization, we compared language network assignments from population and individualized parcellations against task-derived language maps. Individualized language rsFC networks more closely matched same-individual task-based maps than population rsFC networks (g) or other-individual rsFC networks (h). Individualized vs. population rsFC networks, Wilcoxon signed-rank test (n=21): AUC metric, p=5.1×10^−3^; correlation metric, p=1.6×10^−2^. Same-vs. other-individual rsFC networks, permutation test (20000 permutations): AUC metric, p<0.001; correlation metric, p<0.001.

**Extended Data Fig. 17.**
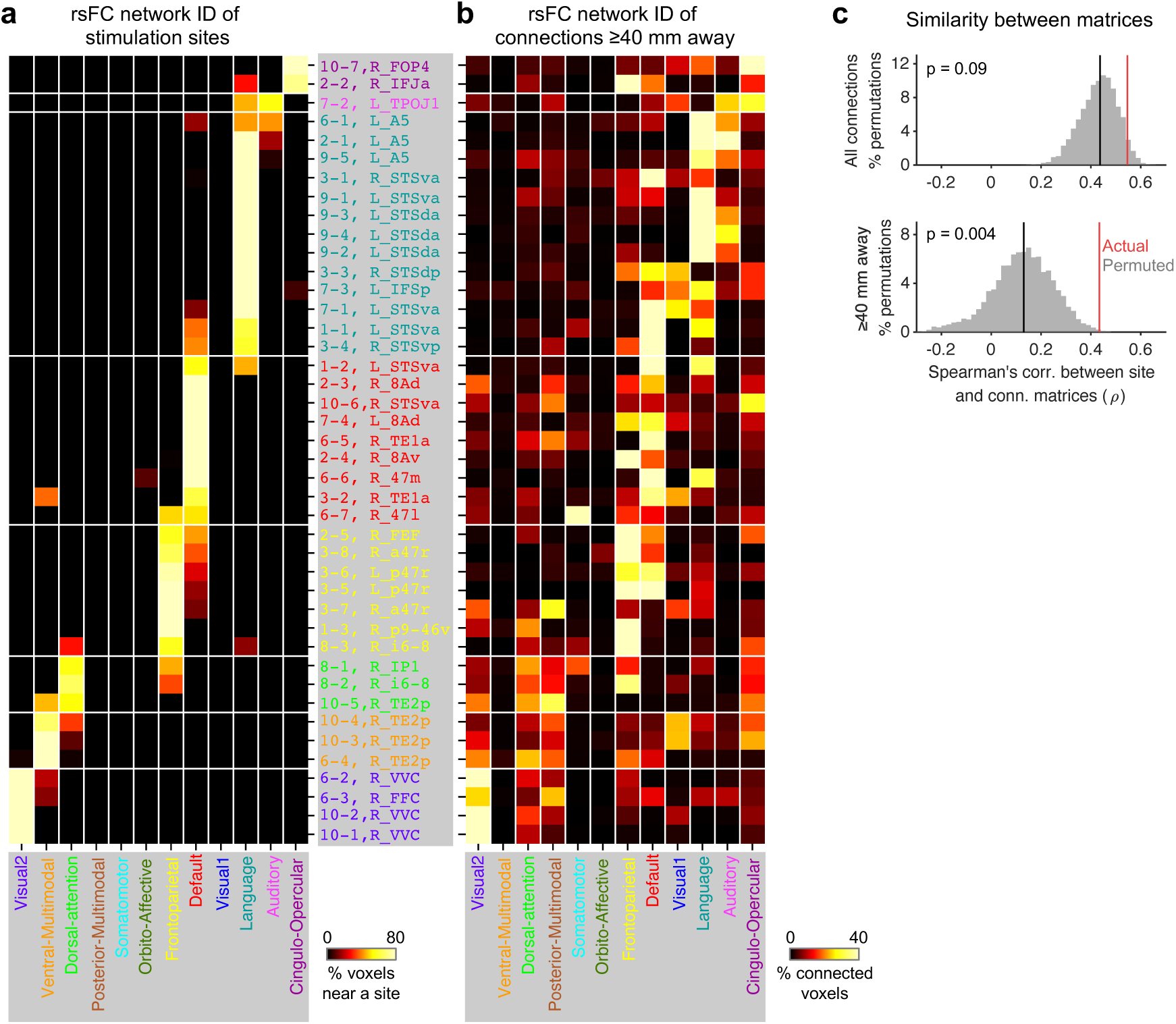
Connectivity within and across rsFC networks. (a,b) Network×site matrices analogous to the contrast×site matrices in Fig. 3a,b, but based on individualized rsFC network assignments instead of task fMRI activations. (a) Percentage of voxels near each stimulation site assigned to each of the twelve rsFC networks. Columns and rows are ordered to highlight the diagonal pattern. (b) Percentage of connected voxels (≥40 mm away) assigned to each network (among all connected voxels ≥40 mm away), with the same ordering as (a). (c) Histogram of SPIN-permuted (20000 permutations) similarity values (Spearman’s ρ) between two matrices in (a) and (b). Here, we jointly rotated and displaced all stimulation sites and their connections on the cortical surface while keeping rsFC networks fixed. Top: using all cortical voxels, excluding those near the site (≤4 mm). Bottom: excluding voxels <40 mm from stimulation sites. Red lines indicate the actual value. Black lines indicate the median of SPIN-permuted values. For all connected voxels, the actual similarity (Spearman’s ρ) was only marginally higher than the permuted value (actual=0.547, median/max=0.437/0.688, two-sided p=0.09). It was common for a large piece of continuous cortex to belong to the same rsFC network; therefore, many connected voxels not too distant from a stimulation site may fall within the same network as the site (more easily than the task-based activation of the same function) after SPIN permutation. When voxels <40 mm from the stimulation sites were excluded, the actual similarity became significantly higher than the permuted value (Fig. 4c, bottom; actual=0.434, median/max=0.130/0.476, p=3.9×10^−3^), suggesting connections are more likely to occur within (than across) rsFC networks.

**Extended Data Table 1.**
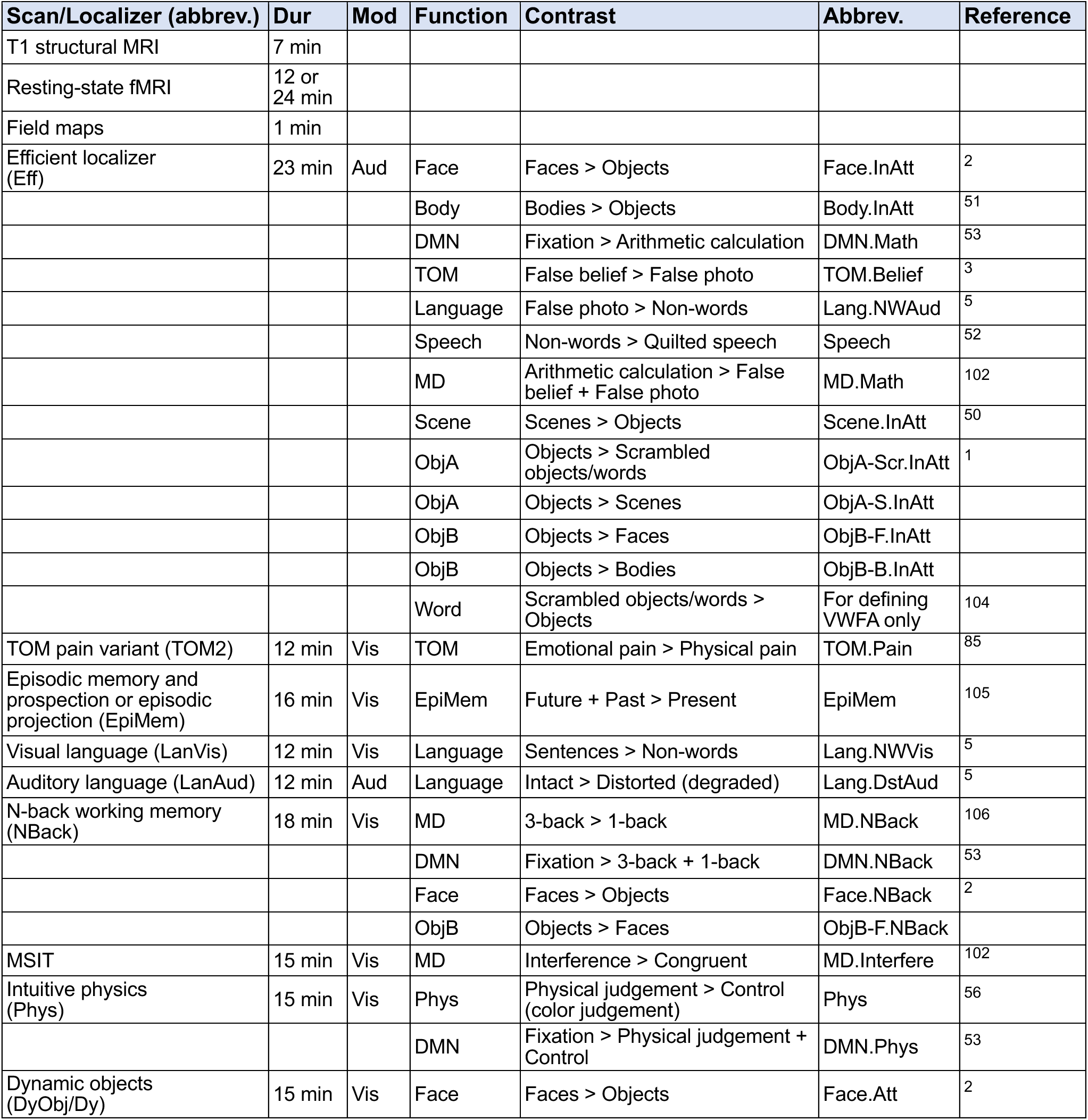
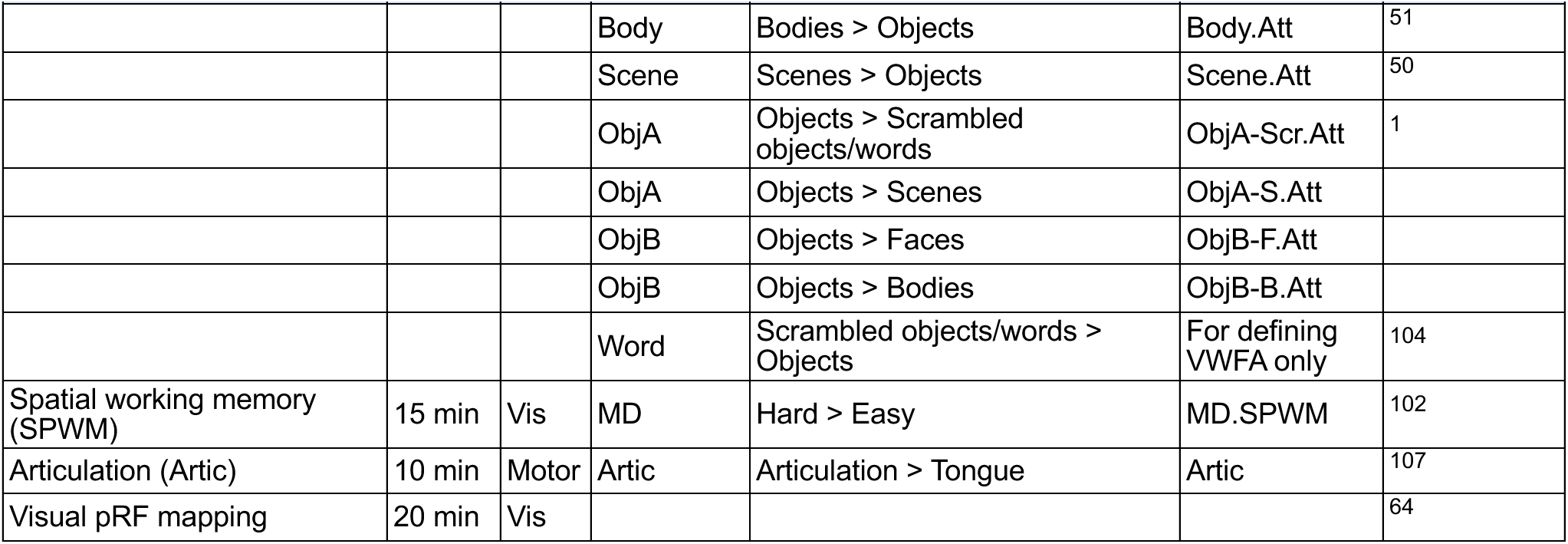
Task-based fMRI localizers and contrasts. The battery included 32 contrasts targeting 13 functions, acquired across multiple task-based localizers in noninvasive MRI sessions. Not all localizers/contrasts were administered to every participant (Extended Data Table 2). Additional object contrasts (objects > faces, objects > bodies, objects > scenes) were included to characterize voxels in the high-level visual cortex that prefer objects over other categories but do not show classic object > scrambled responses. These contrasts were grouped into two functions (ObjA and ObjB) based on the spatial correlation between their cortex-wide activation patterns (Extended Data Fig. 5). We included the word contrasts solely to localize the VWFA region in the left ventral temporal cortex. Because movies of superimposed words and scrambled objects could capture more attention than movies of intact objects, we didn’t use the word contrasts in functional profile analyses. Contrast abbreviations match the labels in Fig. 3a. Localizer abbreviations match the labels in Fig. 4b. Dur, duration; Mod, modality; Aud, auditory; Vis, visual; InAtt, inattention (participants performed auditory tasks in the efficient localizer while viewing visual stimuli passively); Att, attention (visual stimuli were presented without auditory tasks in the dynamic objects localizer).

**Extended Data Table 2.**
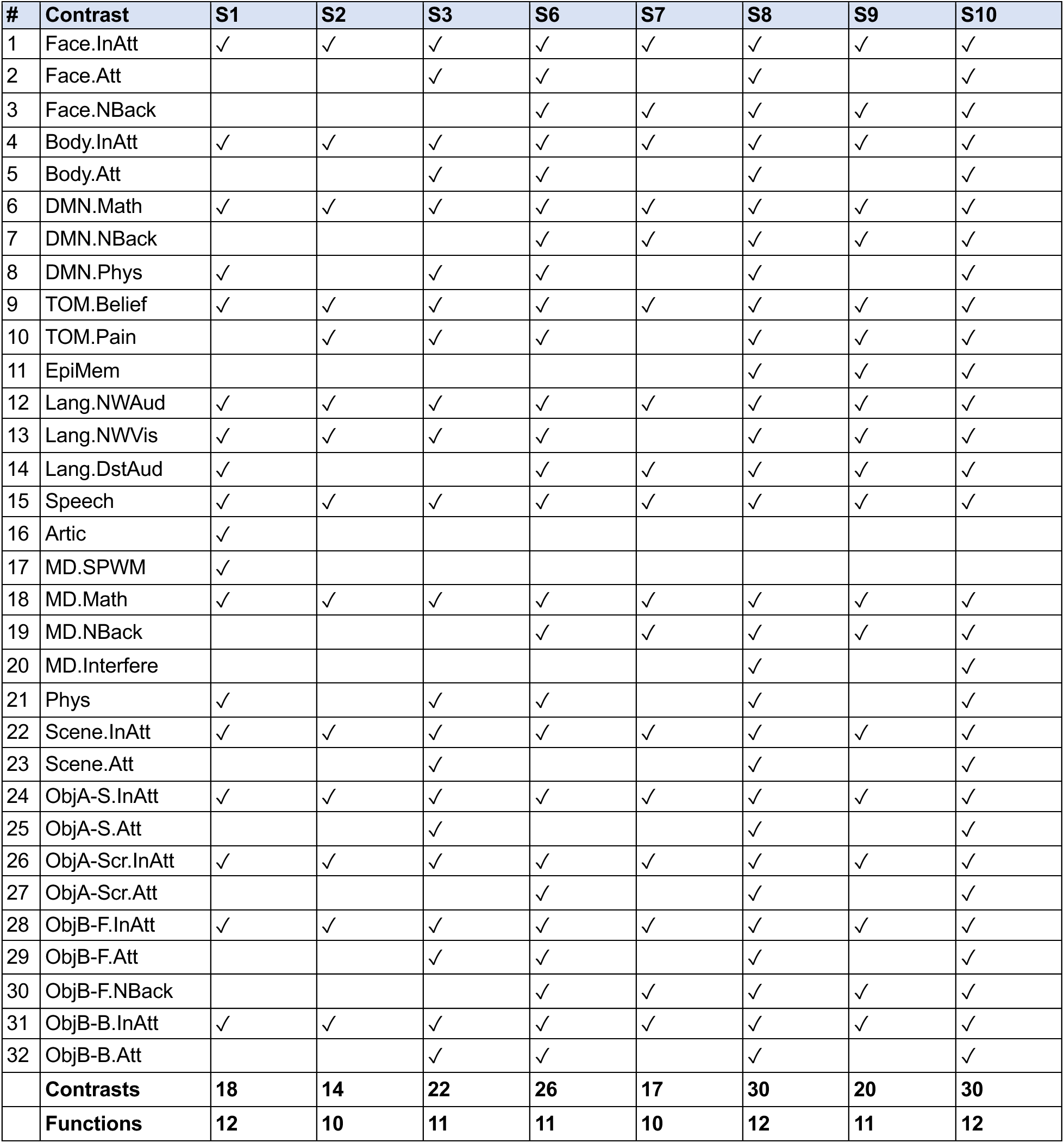
Contrast availability per participant. Check marks indicate contrasts available for each of the eight participants who completed noninvasive MRI sessions.

